# Dual effects of influenza A virus PA-X on suppression of cytokine signaling and MHC I pathway disruption in the human airway epithelium

**DOI:** 10.64898/2026.01.30.702929

**Authors:** Alessandra C. Setaro, Corinne Thomas, Marta M. Gaglia

## Abstract

The mechanisms of immune evasion of influenza A virus are key to its success as an infectious agent. To replicate effectively, influenza A virus must suppress both early innate immune responses, like antiviral type I and III interferons, and adaptive immune responses. The latter is particularly important during infections of animals and humans with pre-existing immunity and explains the continued circulation of influenza A viruses. Here we report how influenza A virus employes a single immunomodulatory protein, the endoribonuclease PA-X, to modulate both arms of the immune responses. To define how influenza A virus uses PA-X to evade immune responses, we characterized its impact on the host response to infection in the infected and bystander cells of the airway epithelium using a 3D ex vivo model and primary cells from multiple human donors. PA- X significantly decreases secretion of multiple cytokines from the airway epithelium, including IFN- λ, inflammatory cytokines, and growth factors related to lung damage, which likely plays a role in its ability to modulate inflammation and lung pathology in vivo. In addition, we discovered that PA-X also decreases surface levels of major histocompatibility complex I (MHC I) on infected cells. This reduction may impact antigen-specific T cells and reduce adaptive immune detection. These dual functions for PA-X highlight how influenza A virus employs active mechanisms to block not only innate immunity but also adaptive immune detection, in addition to tolerating high levels of antigen mutations to escape it.

**Author Summary:** Influenza viruses must evade both innate and adaptive immune responses to infect, replicate, and transmit to new hosts. Through innate immunity, infected cells activate proteins to disrupt viral replication and alert surrounding cells. Through adaptive immunity, antigen-specific T cells recognize specific viral peptides and kill infected cells. We studied how influenza overcomes immune responses. Using a model of human airway tissue, we found a single influenza protein PA-X, can impinge on both innate and adaptive responses. PA-X decreases innate immune signaling between infected and surrounding cells and interferes with the major histocompatibility class I antigen presentation pathway. These dual functions allow PA-X to help the virus continue to circulate and infect new hosts, even in populations with immunity to the flu.

## Introduction

Globally, ∼400,000 people die from influenza A infection and related complications every year^1,2^. The continued burden of influenza A virus is due to its ability to reinfect the same individual multiple times over the course of their life. Indeed, the virus has evolved the ability to outmaneuver host antiviral responses in humans and animals. Manipulation of both innate and adaptive immunity is crucial for successful influenza A virus spread and transmission. Influenza A viruses must evade innate immunity to establish the initial infection. During a primary infection, the adaptive immune system takes over later in infection and contributes to final clearance through the activity of cytotoxic T cells^3,4^. In addition, influenza A viruses often circulate in populations that have been previously exposed to them and have some level of prior immunity. Thus, the ability of influenza A viruses to interfere with adaptive immunity and counteract these memory responses is central to their ability to persist in animals and human populations. Influenza A viruses use a multitude of mechanisms to deal with both arms of the immune responses, which are still being defined in the hope of exploiting them to improve therapeutic approaches and vaccines.

During influenza A infection, the antiviral innate immune response starts in the airway epithelium and then sets the tone for the broader host response. A key cellular response to influenza infection is the release of the antiviral cytokines type I and III interferons (IFNs)^5^, which block influenza infection and communicate the presence of an infection to neighboring cells and immune system cells^6^. Deficiencies in IFN signaling lead to more severe influenza infections in patients^7–9^. However, while IFNs are protective early in influenza infection, excess IFN signaling triggers inflammatory responses that drive severe disease, contributing cytokine storms^10^ and disrupting pulmonary epithelium repair^11^. In addition to IFN signaling during infection, other cytokines shape the host response in the airways. Pro-inflammatory cytokines play key roles in mediating the early influenza A response, and they must strike the right balance to effectively combat infection without causing additional damage to the host^12^. Influenza A virus encodes multiple immunomodulatory proteins that target this early host antiviral signaling and disable communication to neighboring cells^13–15^.

Adaptive immunity is also key for clearance of influenza infection. Mice that have intact innate but impaired adaptive immune responses (RAG2^-/-^ and SCID) are unable to clear influenza infections^16,17^. However, the low vaccine efficacy and the poor adaptive immune responses following illness demonstrate that influenza A viruses can circumvent these responses too ^18,19^. Limited vaccine efficacy is generally attributed to antigen mutations as the virus circulates. However, human challenge studies have shown that people can be re-infected with the same exact influenza A strain as soon as a year following initial infection^19^. These reinfections occurred even in individuals with detectable antigen-specific antibodies to the viral surface antigens hemagglutinin and neuraminidase^19^. This indicates that additional mechanisms are at play to prevent effective adaptive immune responses. A recent study pointed to downregulation of major histocompatibility complex I (MHC I) antigen presentation in infected cells by influenza A virus as a likely strategy of viral evasion of adaptive immunity^20^. A reduction in antigen presentation provides an opportunity for the virus to replicate prior to detection from antigen-specific T cells^21^. In transformed cell lines, Koutsakos et al. reported that influenza A virus decreases surface MHC I levels, and linked this decrease to a change in the total levels of MHC I, rather than a trafficking defect^20^. However, they did not identify the mechanism leading to MHC I downregulation, or determine whether it occurs in the human airway epithelium^20^. Whether viral proteins directly contribute to this mechanism of adaptive immune evasion is unknown. Moreover, while influenza A protein NS1 interferes with the induction of adaptive immunity in the naïve host^22–24^, in general there are limited data for the involvement of influenza A virus proteins in evading the antigen-specific adaptive immune responses.

The RNA endonuclease PA-X is the most recently described influenza A virus immunomodulator, and its function is still unclear^25^. PA-X is an accessory protein produced from the polymerase acid (PA)-encoding segment 3 of the influenza A genome through a +1 ribosomal frameshift (Figure 1A)^25,26^. This frameshift results in a protein with the same N-terminal ribonuclease (RNase) domain as PA and a unique C-terminal domain termed the X-ORF (Figure 1A)^25,26^. At a cellular level, the PA-X RNase activity triggers widespread downregulation of host gene expression during infection, also known as host shutoff^25^. Consistent with its host shutoff function and the general link between host shutoff and immune regulation in many viruses^27^, PA-X generally dampens immune responses *in vivo*^25,28–34^. Infection with PA-X-deficient influenza A strains results in increased expression of immune response genes in the infected airways compared to infection with wild-type (WT) influenza in multiple strain backgrounds and model organisms^25,28–34^. Broad changes in cytokine secretion *in vivo* have also been reported^28,32,34^. However, how PA-X activity alters secretion of cytokines from epithelial cells and how these signals are propagated to uninfected neighboring cells remains unknown. There is some evidence in the literature that PA-X directly downregulates type I and III IFN mRNAs during infection^33,35,36^, but protein-level IFN changes have not been detected consistently^36^.

**Figure 1.**
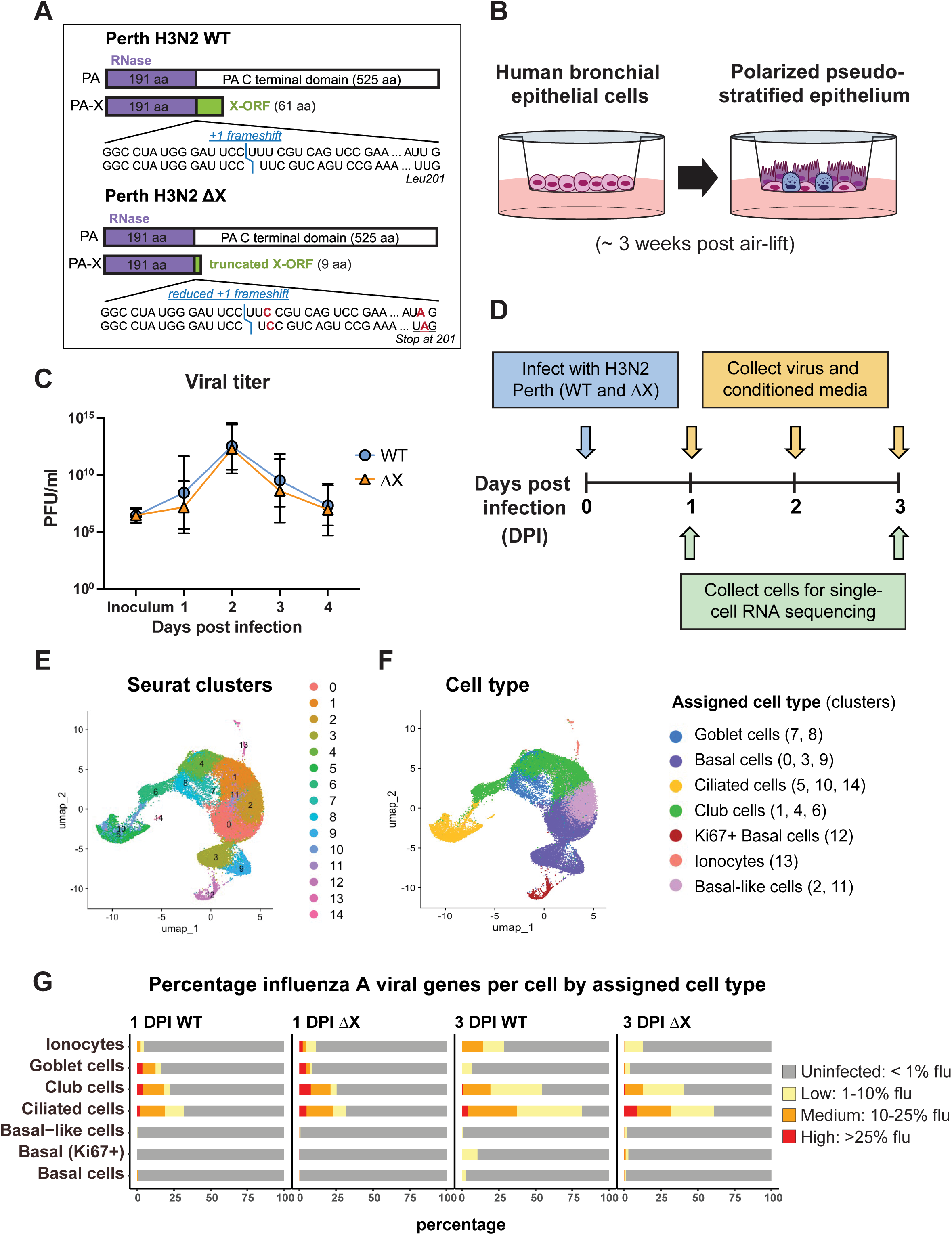
Establishing Perth H3N2 infection kinetics in air-liquid interface (ALI) cultures as a model to study PA-X activity in the airway epithelium. A) Diagram of PA-X production mechanism and mutations in the PA-X defective Perth H3N2 ΔX virus. The position of the PA-X frameshift, identical in 99% of influenza A virus strains, is indicated in blue. The mutated nucleotides that reduce frameshift and generate a premature stop codon at PA-X codon 201 in the ΔX virus are in red. B) Schematic of ALI culture differentiation from primary basal bronchial epithelial cells. C) Virus titer from apical washes of ALI cultures (n = 11, 5 separate donors) measured by Tissue Culture Infectious Dose 50 (TCID50) D) Experimental timeline for infection and sample collection of ALI cultures for all experiments except Calu-3 experiments in Fig. 4E-G. E-G) ALI cultures were infected with Perth H3N2 WT or ΔX at a multiplicity of infection (MOI) of 0.1, or mock infected, and samples were collected as described in Figure 1D. scRNAseq data from Perth H3N2-infected ALI cultures from both donors were analyzed to identify groups of cells with similar expression patterns and to assign cell type. E) Integrated UMAP diagram of all samples, showing the 15 clusters unbiasedly identified by the program Seurat. F) Integrated UMAP diagram of all samples displaying epithelial cell types assigned by ScType. The clusters mapping to each cell type are listed in parentheses. G) Bar graphs with percent influenza A viral reads per cell in each assigned cell type in WT and ΔX infected cultures at 1 and 3 DPI.

Surprisingly, immune modulation by PA-X does not consistently help the virus replicate, a departure from the conventional role of viral immunomodulators. While some *in vivo* studies, especially in avian viruses, have reported that PA-X confers a replication benefit^28,33,37–39^, others show that PA-X has no effect on viral titer^25,32^ or even that it reduces replication^29–31^. In addition, PA-X has variable impacts on disease severity and survival in *in vivo* infection studies in different strains and model organisms^25,28–34,37–39^. Thus, the impact of PA-X during infection is not well understood, and it remains unknown which specific responses are affected by PA-X activity and how they ultimately modulate immune signaling. Nonetheless, the phenotypes reported so far indicate that PA-X has a role in the regulation of immune response that deviates from simply supporting viral replication in the face of early innate immune responses. Moreover, despite its lack of impact on viral replication, PA-X must provide a yet-unknown important benefit to the virus, as it is retained in 99% of influenza A virus strains, as evidenced by the identical frameshifting motif found in these strains as well as the conserved presence of overlapping C-terminal reading frames in segment 3^25,26^.

In this study, we sought to learn how PA-X influences the host antiviral response and circumvents immunity in the context of the complexity of the airway epithelium. The airway epithelium is composed of multiple cell types that play different roles in maintaining barrier function and homeostasis in the respiratory mucosa and in coordinating an infection response, and that have different transcriptional profiles and signaling capacity^40^. The human airway epithelium can be modeled *ex vivo* using air-liquid interface (ALI) cultures^41–43^ (Figure 1B) generated from primary basal stem cells from human donors differentiated by exposure to air^41,42^. The cells in these cultures form a polarized pseudo-stratified epithelium that reflects the complexity of the airway epithelium and contains diverse respiratory epithelial cell types, including basal, ciliated, goblet, and club cells^43,44^. Using this system, which also allowed us to separately analyze infected and bystander cells, we discovered that PA-X has a dual role in thwarting host immune responses: it not only suppresses initial antiviral innate immune signaling at the site of infection but also interferes with protein expression needed for an effective cell-based adaptive immune response. PA-X activity reduced expression and secretion of multiple paracrine signals, including type III IFNs, pro-inflammatory cytokines, and growth factors involved in lung injury. In addition, PA-X activity significantly decreased MHC I in infected cells and slowed MHC I trafficking to the cell surface during infection. Thus, PA-X likely contributes to evasion of antigen-specific immune responses. The dual activity on both innate and adaptive responses points to PA-X as a key player in the continued success of influenza A viruses in the face of host immunity.

## Results

### Influenza A virus replicates robustly in an air-liquid interface (ALI) model of the human epithelium in the presence or absence of PA-X activity

PA-X limits host gene expression during infection in infected cells and broadly alters immune response markers *in vivo*^25,28–34,36–39,45–49^. However, it remains unclear how these two effects are mechanistically connected and which mediators underlie the direct effects of PA-X on immune responses. We thus sought to define PA-X effects on the airway epithelium, the first site of infections. We infected ALI cultures with WT and PA-X deficient (ΔX) viruses in the background of the A/Perth H3N2/16/2009 H3N2 (henceforth Perth H3N2) strain (Figure 1A). Perth H3N2 is a representative of the currently circulating seasonal H3N2 subtype, which is responsible for approximately half of influenza infections in the United States every year^50^. To generate Perth H3N2 ΔX, we combined a mutation that reduces frameshifting events and one that introduce a premature stop codon in the PA-X ORF to ensure that no active PA-X was produced (Figure 1A)^46^. These mutations are silent in the PA reading frame, which prevents confounding changes in PA activity (Figure 1A). We successfully infected ALI cultures with Perth H3N2, resulting in virus production over a 4-day infection time course (Figure 1C). To control for donor-to-donor variability, we performed experiments in cells from multiple young to middle-aged healthy adult males and females (Supplemental Table 1). PA-X mutation did not change viral titers or replication kinetics, as previously reported for other strains of influenza (Figure 1C)^25,32^.

To define the effects of PA-X on the transcriptome of different cell types of the airway epithelium, we collected samples for single-cell RNA sequencing (scRNAseq) at 1 and 3 days post infection (DPI) or mock-infected ALI cultures (Figure 1D). Seurat-based analysis identified 15 cell clusters with distinct gene expression patterns within the cultures (Figure S1A). Using ScType, a program that assigns the most likely cell identity to each cluster based on the expression of marker genes from a lung cell gene database^51,52^, we determined that the clusters recapitulated the major cell types expected in the airway epithelium – goblet, basal, ciliated, club cells, and ionocytes (Figure 1E-F, S1A). Moreover, our ALIs display tight junctions, cilia, and mucus production, recapitulating the microphysiology of a healthy respiratory epithelium (Figure S2A), and can be infected by Perth H3N2 (Figure S2B). We verified the cell type assignment by confirming that appropriate key markers were expressed in corresponding clusters, which also allowed us to further classify some epithelial subtypes based on known markers and pseudo time analysis (Figure S1A-B). We also confirmed that the proportion of basal, ciliated, club, and goblet cells was similar between the ScType-based assignment and classical flow cytometry analysis of established epithelial cell type markers (Figure S2C-D)^53^. Having determined that the ALI cultures recapitulated the complexity of the human airway epithelium, we examined which cells were infected with Perth H3N2 and whether PA-X influenced the tropism. We quantified the percentage of influenza A reads in different cell types and clusters. Perth H3N2 readily infected ciliated and club cells, and goblet cells to a lower extent (Figure S3A-B), similar to what has been previously reported for A/Udorn/307/72 H3N2^54^ and 2009 pandemic H1N1 strains^44,55^. We also detected changes in the proportion of different cell types during infection, which in part reflected the death of infected cells at 3 DPI (Figure S2D). However, there were no significant differences in target cell type in WT vs. ΔX-infected cultures (Figure 1G), and the proportion of different cell types also changed in a similar manner regardless of PA-X activity (Figure S2D).

Taken together, our initial characterization shows that in our model of the bronchial epithelium, PA-X has no apparent impact on viral titers or cell tropism. Therefore, this system allows us to dissect PA-X-dependent differences in virus-host interactions without the confound of altered replication.

### The magnitude and quality of the transcriptional response to influenza A virus infection and PA-X activity vary depending on the cell subtype

Since epithelial cell populations have different roles in tissue homeostasis and infection responses, they are likely to display different transcriptional responses to infection and PA-X activity. To define these responses, we analyzed differentially expressed genes (DEGs) in each cluster using Seurat’s FindAllMarkers function^56^. Many DEGs were detected between Perth H3N2 WT and mock infections (Table 1A), illustrating the large transcriptional changes caused by influenza A viral infection of the airway epithelium. Surprisingly, several cell types – ionocytes (cluster 13), deuterosomal ciliated cells (cluster 14), and proliferating Ki67+ basal cells (cluster 12) – exhibited limited transcriptional changes in response to infection, particularly at 1 DPI.

**Table 1.**
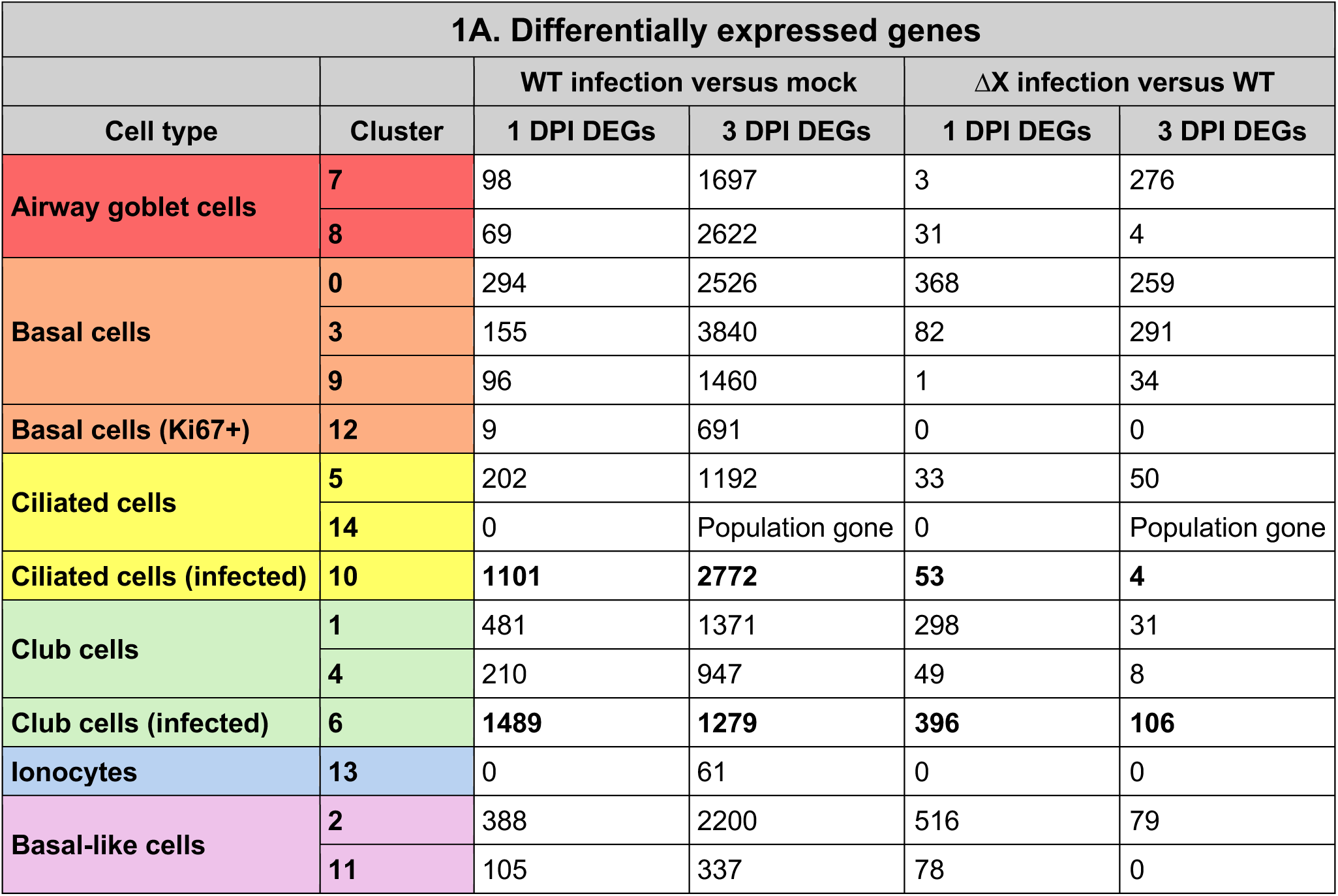
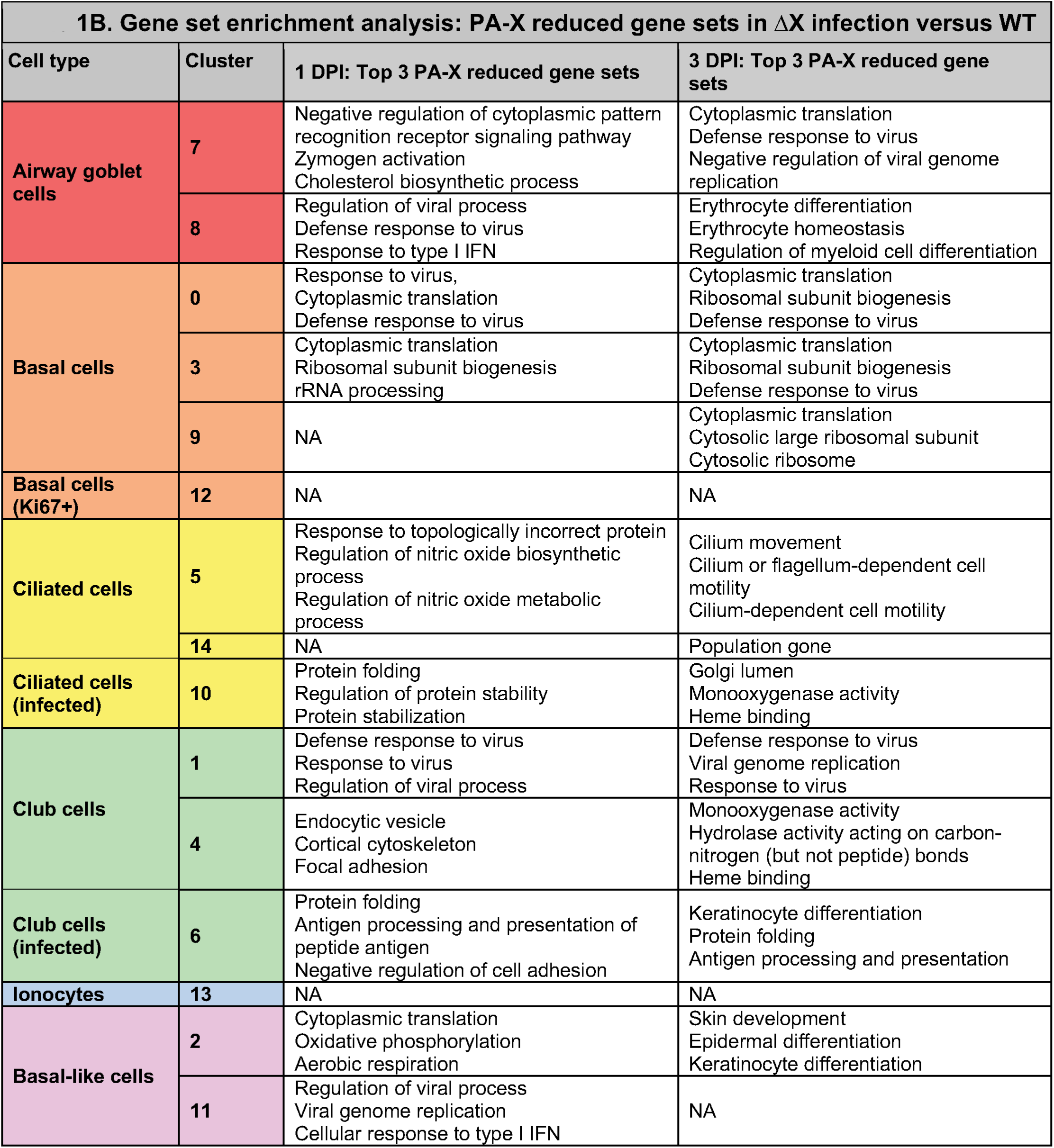
Different subtypes of cells respond differently to influenza infection and PA-X activity. ScRNAseq data from ALI cultures infected as shown in Figure 1D were analyzed to identify infection and PA-X-driven changes in gene expression. Differentially expressed genes (DEGs) were identified using FindAllMarkers with a cutoff of |log2FC| > 0.25 p-adjusted < 0.05, and enriched gene set cutoffs were p-adjusted < 0.05 (n = 2, Donor 1 and Donor 2). A-B) Number of DEGs per cluster/cell type in WT/mock infections (A) and ΔX/WT infections (B). C) Gene set enrichment analysis was carried out on PA-X-decreased DEGs by cluster/cell type. The top 3 enriched gene sets are listed for each cluster/timepoint.

We used gene set enrichment analysis^57–59^ to determine which biological processes were regulated by influenza A virus infection. Gene sets related to microtubule formation were enriched in upregulated genes in many cell types (Table S3), which is consistent with influenza A virus using tunneling nanotubes to traffic viral ribonucleoproteins (vRNPs) within and between cells^60–62^. Cell cycle-related functions (for example: mitotic nuclear division, mitotic sister chromatid segregation) were downregulated in infected cells, consistent with an increased proportion of cells arrested in G1 (Figure S4A-B), likely due to a known activity of the influenza A protein NS1^63,64^. These overall changes in infected cells illustrate how the virus immediately begins to reshape the host cell environment for its own benefit. While there was an increase in cytokine-related genes, specifically response to interleukin 1 and regulation of leukocyte migration, surprisingly there was no enrichment of interferon signaling related gene sets in WT vs. mock infection until 3 DPI (Table S3).

To determine how PA-X activity altered gene expression in different cell types during infection, we analyzed DEGs in Perth H3N2 ΔX vs. WT-infected cultures (Table 1A). At 1 DPI, PA-X activity caused the largest differences in gene expression in club cells (clusters 1, 6), basal-like cells (cluster 2), and basal cells (cluster 0) (Table 1A). By 3 DPI, basal (cluster 0,3) and goblet cells (cluster 7) had the most PA-X dependent DEGs. Similar to the WT vs. mock comparison, there were no significant gene expression changes in response to PA-X activity (ΔX vs. WT) in ionocytes (cluster 13), deuterosomal ciliated cells (cluster 14), and proliferating basal Ki67+ cells (cluster 12) (Table 1A). We used gene set enrichment analysis^57–59^ to investigate PA-X-downregulated gene sets, as PA-X is an endoribonuclease that reduces host gene expression^25,45,46,65^. In most clusters, PA-X activity decreased expression of genes relating to viral defense response and IFN response, indicating that it was at least in part responsible for the lack of a clear early antiviral signature in WT vs. mock infected cells. This result further supports the model that PA-X’s main role is to suppress host antiviral responses. There were some additional notable and unexpected results. In ciliated cluster 5, there was a reduction in stress-related gene sets, including regulation of nitric oxide (NO) formation, at 1 DPI and in genes related to cilia movement at 3 DPI. Cilia motility is crucial to mucociliary clearance and NO increases ciliary beat frequency to efficiently clear respiratory pathogens^66,67^. Also, in infected club (clusters 6) and ciliated (cluster 10) cells, there was a PA-X-dependent decrease in genes involved in antigen processing and presentation at both 1 and 3 DPI (Table 1B). A previous study showed that influenza A virus infection decreases major histocompatibility (MHC) class I, but the mechanism for this has not yet been established^20^.

Taken together, these results highlight the heterogeneity of the response to infection by airway cell types. While the PA-X-dependent changes in antiviral response provide further support for the immunomodulatory role for PA-X, the scRNAseq also revealed potential novel functions for PA-X, including regulation of antigen presentation.

### PA-X activity blunts the type I and III IFN response in human bronchial epithelial cell ALIs

Type I and III IFNs are major players in the antiviral response. In other studies, PA-X was reported to reduce antiviral type I and III interferon (IFN) mRNAs during infection in model organisms and transformed cell lines^33,46,49^. However, a previous study indicated that PA-X-dependent decrease in IFN mRNA levels did not lead to changes in IFN secretion during *in vivo* infection^36^. Thus, the effect of PA-X activity on type I and III IFN secretion during infection remains unclear. Our results indicate PA-X activity decreased expression of genes in the IFN response gene set during infection (Table 1B). To further investigate how PA-X activity altered IFN expression, secretion, and responses throughout infection in a human epithelium-like setting, we compared expression of type I and III IFN genes in all clusters, separated by infection status (infected = viral gene positive; bystander = viral gene negative), viral strain, and time point (Figure 2A). Type I and III IFN genes were expressed in infected cells, but not in uninfected bystander cells (Figure 2A). This is consistent with IFNs being produced by infected cells in response to sensing of viral RNAs^5^. Moreover, their levels were consistently elevated in ΔX-infected cells compared to WT infection at 1 but not 3 DPI (Figure 2A). As expected based on the infected cell types, IFNs were expressed predominantly by ciliated and club cells (Figure 2B). Since previous studies have reported PA-X-dependent changes in IFN mRNAs but not protein^36^, we tested whether these transcriptional changes led to changes in IFN secretion into the ALI basal media. PA-X activity decreased secreted IFN-λ at 2 and 3 DPI, with a nearly two-fold increase in IFN-λ in ΔX vs. WT-infected ALIs at 2 DPI (Figure 2C). As previously reported in the infected airway epithelium^68,69^, we found that IFN-β was produced at much lower levels (approximately 1000x less) compared to IFN-λ, and PA-X did not alter secreted IFN-β levels (Figure S5).

**Figure 2.**
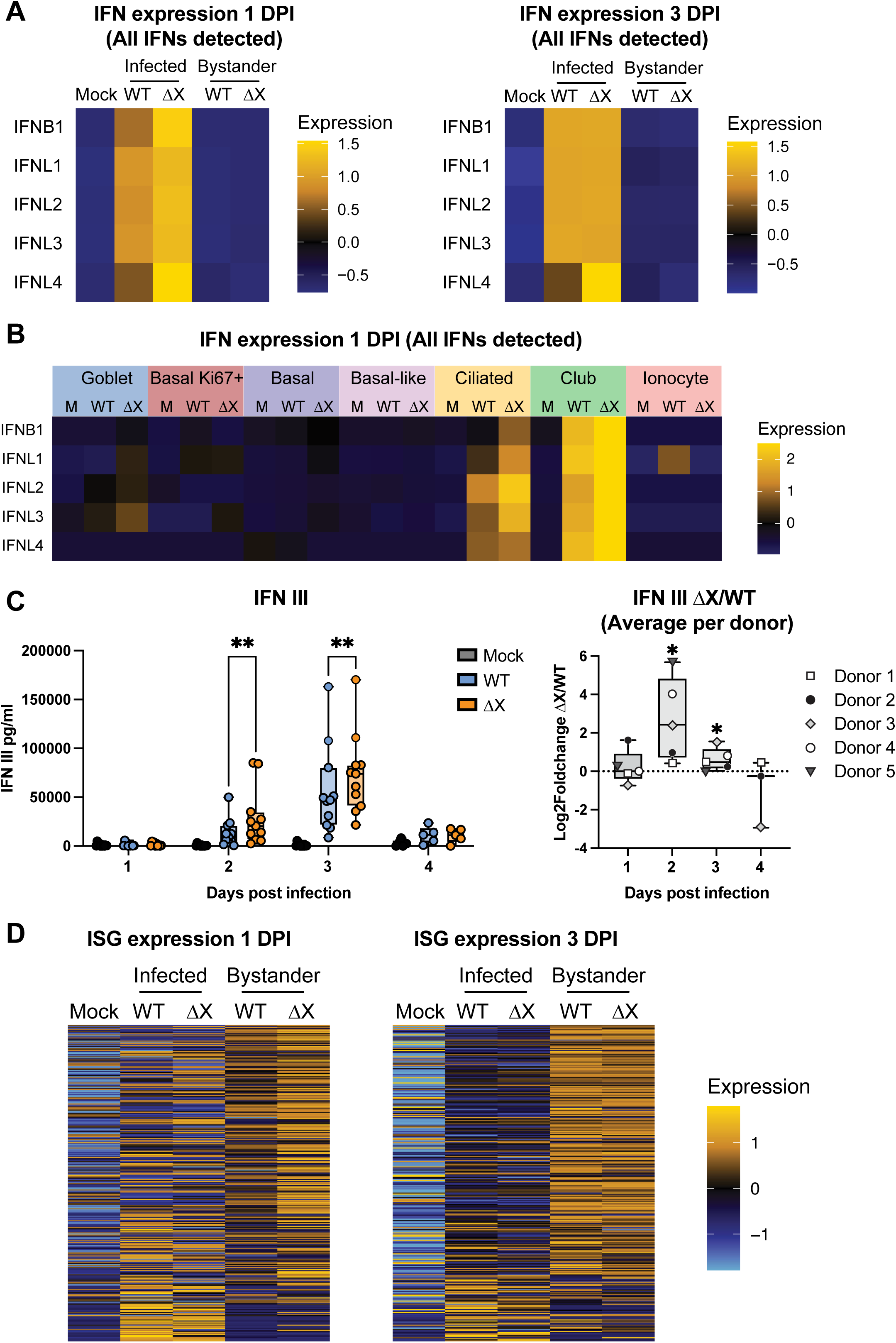
PA-X activity blunts the type I and III IFN response in ALI cultures. ALI cultures were infected and samples collected as described in Figure 1D. The scRNAseq data were analyzed for changes in type I and III IFN production and responses (n = 2, Donors 1 and 2). Basal medium was used to measure secreted IFN-λ (n = 5-11 independent experiments, 5 donors, Donors 1-5). A) Heatmap showing average expression of type I and III IFNs in the scRNAseq data at 1 and 3 days post infection (DPI), separated by the infection status of the cells (infected: ≥ 1% influenza A virus reads; bystander: <1% influenza A virus reads). B) Heatmap showing average expression of type I and III IFNs at 1 DPI in different cell types (assigned with ScType). C) IFN-λ in the basal media was detected using HEK-Blue IFN-λ cells reporter cells. The IFN-λ concentration and log2 fold-change in ΔX vs. WT infection are plotted. ns = p > 0.05, *= p ≤ 0.05, **= p ≤ 0.01, ***= p ≤ 0.001, two-way ANOVA with Šidák correction (IFN-λ concentration) or ratio T-test (Log2 fold-change). D) Heatmap showing average expression of ISGs at 1 and 3 DPI, separated by the infection status of the cells. The list of ISG was based on Schoggins et al. 2011^140^.

We also compared IFN-stimulated gene (ISG) expression in each infection condition to assess responses to IFN. We saw greater ISG expression in uninfected bystander cells compared to infected cells in WT and ΔX-infected cells (Figure 2D). This is likely due to suppression of gene expressions and/or ISGs in infected cells by other influenza proteins like non-structural protein 1 (NS1)^70^. Overall, PA-X activity broadly decreased ISG expression in bystander cells at 1 DPI (Figure 2D), again indicating that PA-X suppresses secretion of type III IFN from infected cells. However, the ISG transcriptional differences appeared prior to detectable changes in IFN secretion at the protein level, suggesting that our protein-based assays were not sensitive enough to detect alterations in IFN secretion at this time point. Despite the continued IFN secretion differences at 2 and 3 DPI, the ISG downregulation was transitory, as there was no net PA-X dependent difference on ISG expression in infected or bystander cells by 3 DPI (Figure 2D). Moreover, there was no PA-X-dependent difference in viral titer throughout a 4-day infection course (Figure 1C). Together, these results demonstrate that PA-X activity reduces type III IFN secretion from airway epithelium during infection, with a greater effect soon after infection, but that this decrease in IFN signaling is not sufficient to affect influenza A viral replication. Nonetheless, these changes in IFN signaling may still contribute to increased inflammatory signaling, immune cell recruitment, and lung injury *in vivo*.

### PA-X limits secretion of pro-inflammatory and lung injury-associated cytokines

Appropriate inflammatory cytokine signaling responses are necessary to combat influenza A viral infections. Patients with single nucleotide polymorphisms in genes encoding pattern recognition receptors^71–76^, pro-inflammatory cytokines^77–82^, and other proteins involved in innate immune signaling^83^ are more susceptible to influenza infections. However, excess production of inflammatory cytokines can result in inflammation and damage during infections^10^. To dissect the full complement of cytokine signals that were controlled by PA-X in the human airway epithelium, we used the program Cytokine Signaling Analyzer (CytoSig). CytoSig uses transcriptional profiles of cells receiving the cytokine signal to predict secreted cytokines^84^. We ran CytoSig on expression data from uninfected bystander cells only, as multiple viral proteins diminish the infected cell’s capacity for host gene transcription, which could confound the analysis^85^. CytoSig analysis predicted changes in the secretion of many proteins between Perth H3N2 WT and ΔX infections at both timepoints. We narrowed down this list further based on detection of their mRNA in our scRNAseq data and availability of Luminex reagents to select proteins for experimental testing. Figure 3A shows the CytoSig prediction for pro-inflammatory cytokines and growth factors we selected for analysis. Pro-inflammatory cytokines were predicted to have elevated secretion in ΔX vs. WT, while growth factors like EGF were predicted to have reduced secretion (Figure 3A). In addition to the predicted cytokines, we also tested CXCL1 and CCL20, which are decreased by PA-X in *in vivo* mouse infections^29,37^ and IGFBP-3, VEGFC, and uteroglobin, which have roles in pathways that were altered based on the CytoSig or GSEA analyses. At 3 DPI, PA-X significantly reduced secretion of the pro-inflammatory G-CSF, IL-6, and IL-1β in the basal media of ALI cultures (Figure 3B-C, S6A). In addition, we also discovered that PA-X significantly reduced secretion of two proteins associated with ARDS and pulmonary fibrosis, IGFBP-3 and FGF basic (Figure 3B-C, S6A)^86–90^. While the detected changes in FGF basic were not dramatic, a clinical study found a slight but significant increase in FGF basic in the bronchoalveolar lavage fluid of patients with chronic beryllium disease and idiopathic pulmonary fibrosis, suggesting that even slight differences are reflective of changes in disease status^88^. Interestingly, none of the signaling molecules tested were higher in Perth H3N2 WT compared to ΔX infection despite the CytoSig results (Figure 3A, S6A-B). PA-X activity thus decreases secretion of multiple cytokines with roles in inflammation and fibrosis during infection, extending the effect of PA-X on innate immune signaling beyond IFN regulation.

**Figure 3.**
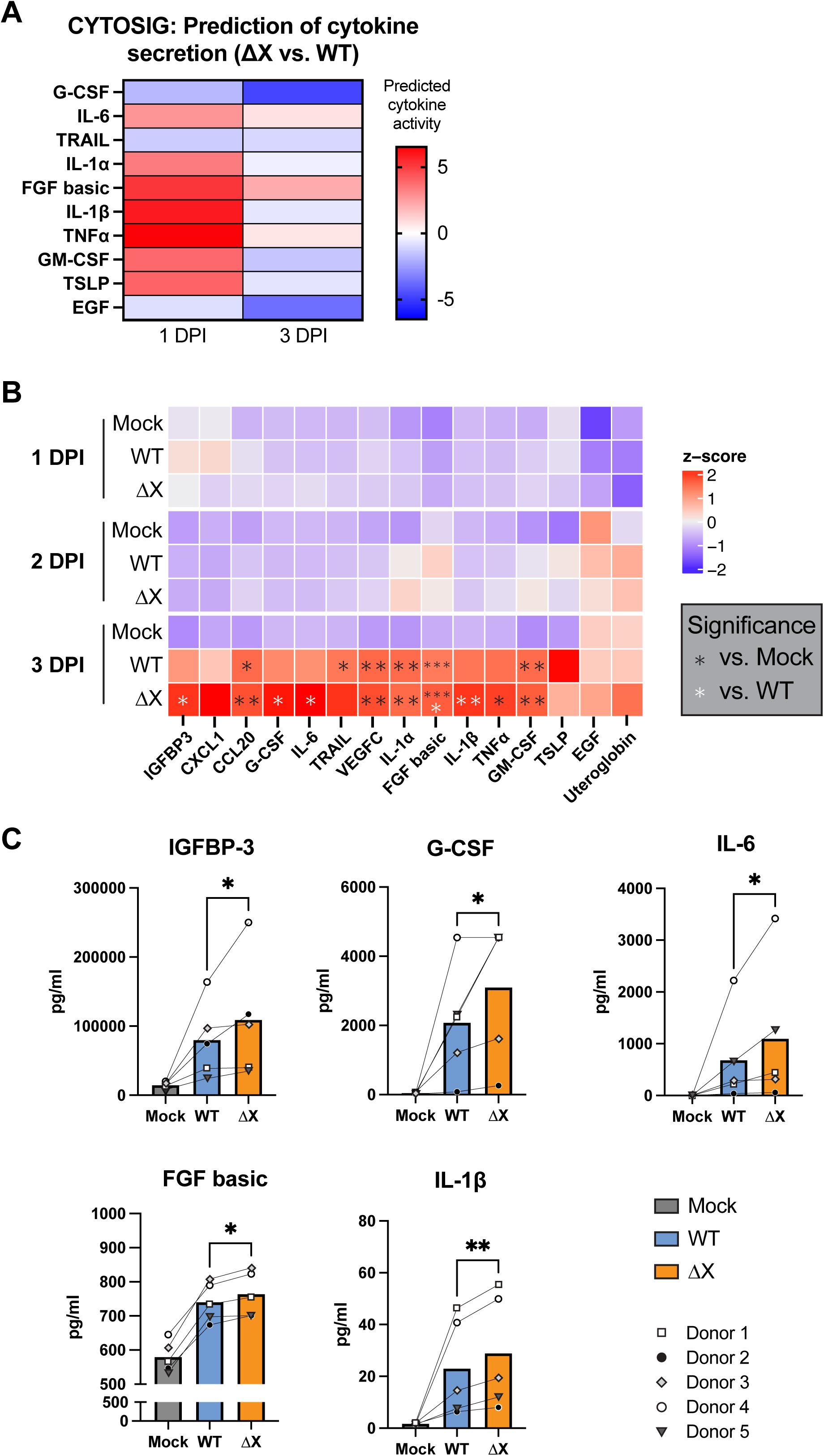
PA-X limits secretion of pro-inflammatory and lung injury-associated cytokines. ALI cultures were infected and samples collected as described in Figure 1D. The scRNAseq data were analyzed using CytoSig for predicted cytokine responses (n = 2, Donors 1 and 2). Basal medium was used to measure secreted IFN-λ (n = 5-11 independent experiments, 5 donors, Donors 1-5). A) Heatmap showing predicted activity (and consequently production) of select targets at 1 and 3 days post infection (DPI) from bystander uninfected cells (<1% influenza A virus reads). B-C) Luminex measurements of cytokine in basal conditioned media, plotted as a heatmap showing mean-centered z-score of the protein concentration for each target, or bar graphs for cytokines that were significantly altered by PA-X activity at 3 DPI. In B, black asterisks correspond to significance vs. mock (two-way ANOVA) and white asterisks correspond to significance vs. WT (ratio t-test). In C, comparisons of ΔX vs. WT were analyzed using a ratio T-test. ns = p > 0.05, *= p ≤ 0.05, **= p ≤ 0.01, ***= p ≤ 0.001.

### PA-X reduces MHC I expression in infected cells

MHC class I presentation is critical to alert the immune system to virally infected cells. To evade antigen-specific immune response, many viruses disrupt the process of presentation of intracellular peptides by multiple mechanisms, including downregulation of surface MHC I levels, the complex that presents peptides to cytotoxic T cells^20,21,91–97^. Recently, Koutsakos et al. reported that influenza viruses decrease MHC I during infection of transformed cells^20^. However, the mechanism of MHC I downregulation by influenza A virus is still unknown. Interestingly, we detected a PA-X-dependent decrease in the expression of antigen processing and presentation genes, especially in club and ciliated cells from clusters 6 and 10, respectively, which include most of the infected cells (Table 1). In addition, an unbiased analysis of functional gene sets^57,58^ differentially expressed in all infected cells due to PA-X activity revealed antigen processing and presentation as one of the top PA-X-downregulated gene sets at both 1 and 3 DPI (Figure 4A). At the single gene level, PA-X activity prevented infection-driven upregulation of MHC I complex components in both infected and uninfected bystander cells, including the genes for the heavy chain, HLA-A, HLA-B, HLA-C, and the light chain B2M (Figure 4B)^98^. PA-X activity also decreased expression of genes involved in antigen processing, such as Transporter Associated with Antigen Processing 1 (*TAP1*)^99–101^ and tapasin (*TAPBP*)^102^, and genes essential for peptide loading, including calnexin (*CANX*)^103^ and calreticulin (*CALR*)^102^ (Figure 4B). PA-X also decreased expression of genes that encode proteasome activator PA28 (*PSME1* and *PSME2*), which plays an important role in the production of MHC I-binding peptides^104^. Additionally, PA-X activity decreases expression of *HSP90AA1 and HSP90AB1*, which encode heat-shock protein 90 (HSP90), a chaperone protein required for MHC I antigen presentation^105^. It is likely that PA-X directly targets several of these mRNAs for degradation. In a previous study of PA-X-dependent RNA cleavage in A549 cells, we detected cleavage sites in 6 of the 21 significantly downregulated genes listed in Figure 4B in cells infected with WT vs. PA-X deficient A/Puerto Rico/8/1934 H1N1 (Figure S7A-B)^65^. To verify that the H3N2 Perth PA-X variant can also cleave these transcripts, we used 5’ RACE (Figure S7A) to detect the cleavage fragments of *CALR* and *HSP90AA1* in cells expressing H3N2 Perth PA-X. Indeed, we detected cleavage fragments for these genes that originated at the same sites as the ones detected in the prior study (Figure S7C-E). While we cannot mutate the catalytic active site of PA-X in the context of the virus, in this experiment we were able to compare wild-type and catalytic mutant (D108A)^106^ PA-X, and confirm that the cleavages were due to the catalytic activity of PA-X.

**Figure 4.**
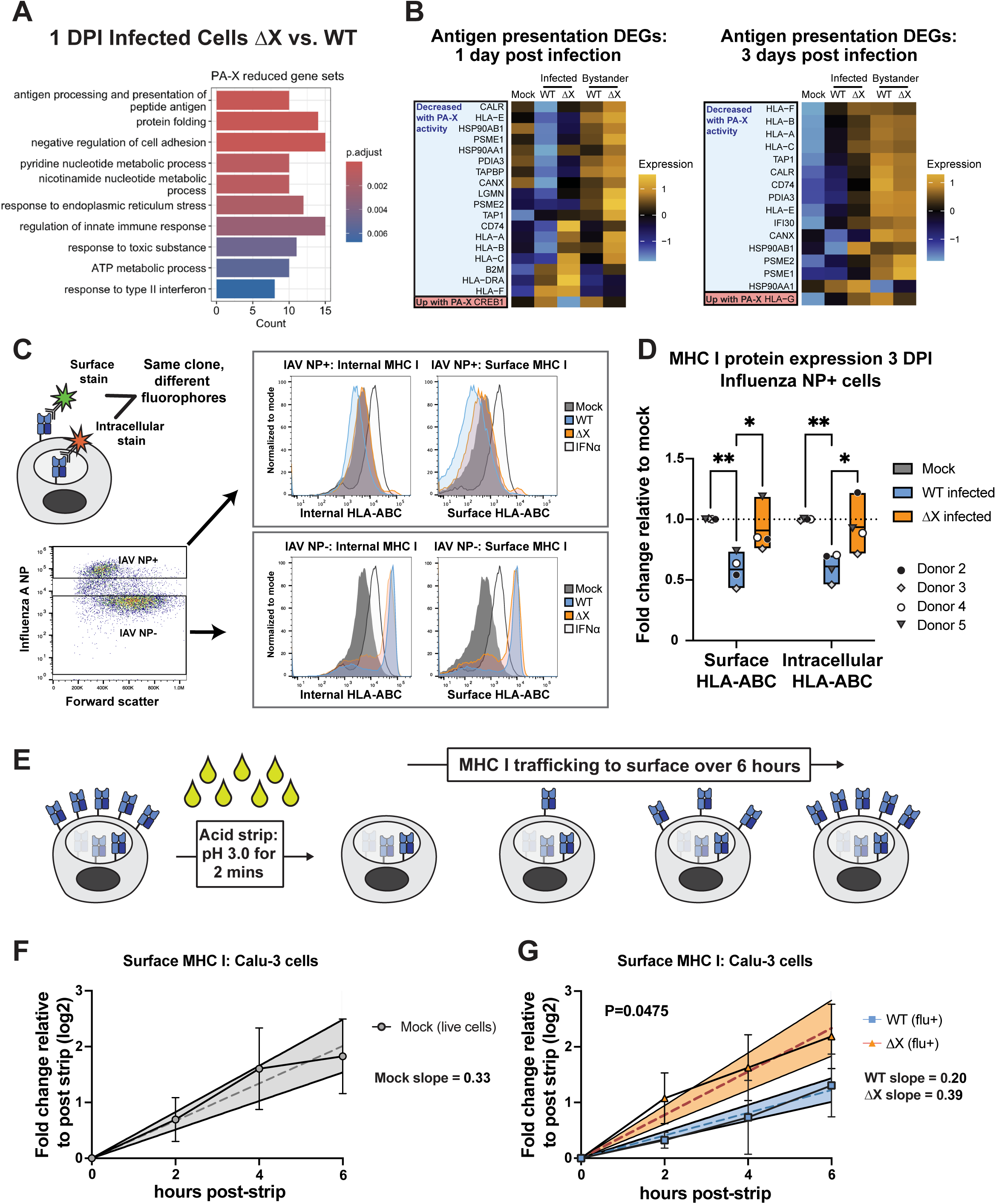
PA-X activity reduces expression of antigen processing and presentation genes and MHC I protein levels in infected cells. A-B) Analysis of scRNAseq data from ALI cultures infected as shown in Figure 1D (n = 2, Donors 1 and 2). A) Top ten gene sets identified by gene set enrichment analysis of differentially expressed genes (DEGs) in infected cells (≥1% influenza A virus reads) at 1 day post infection (DPI). DEGs were identified using FindAllMarkers with a cutoff of |log2FC| > 0.25 p-adjusted < 0.05, and enriched gene set cutoffs were p-adjusted < 0.05. B) Heatmaps of DEGs in the antigen processing and presentation gene set (GO:0019882) at 1 and 3 DPI, separated by by the infection status of the cells (infected: ≥1% influenza A virus reads; bystanders: <1% influenza A virus reads). C-D) ALI cultures were infected as shown in Figure 1D or pre-treated with IFN-α at 2 ng/µL for 1 hour. At 3 DPI, cells were stained for major histocompatibility complex I (MHC I) and influenza A virus nucleoprotein (IAV NP). C) Diagram of the staining method, gating strategy for NP+ infected and NP- bystander cells (after gating for live, single cells), and representative histogram results for MHC I HLA-ABC surface and intracellular expression. The same anti-HLA-ABC coupled to different fluorophores was used before and after permeabilization to detect surface and intracellular MHC I, respectively. D) HLA-ABC median fluorescence intensity at 3 DPI from flow cytometry analysis plotted as fold-change relative to mock-infected cultures (n = 4, 4 separate donors, Donors 2-5). Two-way ANOVA, followed by Šídák’s multiple comparisons test, ns = p > 0.05, *= p ≤ 0.05, **= p ≤ 0.01, ***= p ≤ 0.001. E-G) Calu-3 cells were mock-infected or infected with Perth H3N2 WT or ΔX at an MOI of 0.1 for 18 hours. Surface MHC I was stripped using a citric acid buffer, then cells were collected at multiple timepoints over the course of 6 hours. Surface MHC I was measured by flow cytometry. E) Schematic of acid strip time course and MHC I surface trafficking. F-G) Log2 fold change of surface MHC I median fluorescence intensity in all live mock-infected cells (F) or influenza A virus NP-positive WT and ΔX-infected cells (G). Linear regression was performed for each condition, and the p value calculated to compare Perth H3N2 WT and ΔX NP+ cells. Dashed lines = the line of best fit, shaded regions = 95% confidence interval.

To test whether downregulation of MHC I mRNAs altered MHC I protein levels during infection, we used a flow cytometry assay^20^ to detect intracellular and surface MHC I in infected and bystander cells (Figure 4C). Surface MHC I levels are particularly important, as only peptide-loaded MHC I traffic to the surface, and lower surface levels lead to a reduction in antigen presentation. We fixed single-cell suspensions from ALI cultures and stained with anti-HLA-ABC antibody, which binds to conserved epitopes of these HLAs, to detect the complexes on the cell surface. We then permeabilized the cells to stain for internal HLA-ABC, using the same antibody coupled with a different fluorophore (Figure 4C). We also separated infected from bystander cells by staining for influenza nucleoprotein (NP) (Figure 4C, S8A). IFN-α treatment was used as a positive control, as it increases HLA-ABC surface and intracellular expression in primary human cells^107,108^ (Figure 4C). Perth H3N2 WT infection significantly decreased both surface and intracellular MHC I in primary epithelial cells at 3 DPI relative to mock infection, in agreement with previous data in transformed cells (Figure 4D). In contrast, cells infected with the Perth H3N2 ΔX virus had similar surface and intracellular MHC I levels to mock-infected cells (Figure 4D, S8A-B). This indicates that PA-X decreases the level of MHC I in infected cells, which in turn reduces surface levels of this complex. In contrast to infected cells, MHC I levels of uninfected bystander cells increased after infection, and this upregulation was similar between infection with Perth H3N2 WT and Perth H3N2 ΔX (Figure 4C), despite the PA-X-dependent transcriptional reduction seen in bystander cells (Figure 4B). To investigate whether PA-X reduction in MHC I is conserved in different viral strains, we infected ALI cultures with A/California/04/2009 (Cal) H1N1 and found that Cal H1N1 PA-X also shows a trend of reduction in MHC I levels (Figure S8C-D). While the reduction we found was striking and consistent with the previous study in A549 cells, we wondered whether it appeared soon enough to functionally affect antigen presentation. We did not detect a change in MHC I levels at 1 DPI in ALIs infected with Perth H3N2, but the percentage of NP+ cells was low, which may have masked an effect by reducing the discrimination between infected and uninfected cells. A previous study in mouse dendritic cells showed peptides being displayed as early as 3.5 hours post infection with the A/Puerto Rico/8/1934 H1N1^109^. However, our assays were carried in human epithelial cells with a seasonal H3N2 strain, which have different kinetics of viral replication and will likely have different timing of antigen display. Since steady state surface level changes in MHC I are likely the result of changes in trafficking, we also tested how PA-X impacts the rate of MHC I trafficking to the cell surface as a more sensitive readout that could reveal defects earlier. A reduced rate of MHC I trafficking would result in reduced presentation of peptides from newly synthesized proteins, including viral proteins, even before total surface levels are reduced. To measure MHC I trafficking, we used an established acid strip method to remove MHC I from the cell surface of Calu-3 cells, a lung adenocarcinoma cell line, and monitored recovery of surface MHC I levels over time by flow cytometry (Figure 4E)^20,110^. We used Calu-3 cells for these experiments because surface acid strips must be performed on viable, single cell suspensions and cannot be performed on the 3D ALI cultures^110,111^. As previous study showed that peak antigen presentation occurred at 12-24 hours in infections of Calu-3 cells with seasonal H3N2s^112^, we stripped surface MHCI at 18 hours post infection and measured recovery of surface MHC I levels over from 18 to 24 hours post infection (0-6 hours post strip, Figure S8F). 18 hours was also the time when we started detecting viral nucleoprotein by flow cytometry in these cells. In mock-infected Calu-3 cells, surface MHC I median fluorescence intensity doubled approximately 3 hours after acid strip (Figure 4F). Importantly, Perth H3N2 WT infection halved the rate of surface MHC I recovery (Figure 4G). In contrast, in cells infected with Perth H3N2 ΔX, MHC I was trafficked to the cell surface at a rate similar to that of mock-infected cells, and significantly different from Perth H3N2 WT infection (Figure 4F-G). The effect of PA-X was largely cell-intrinsic, as the rate of MHC I trafficking to the surface was not significantly different in uninfected bystander cells in ALI cultures infected with Perth H3N2 WT vs. ΔX (Figure 4F and S8H). Taken together, these data identify PA-X as the viral protein responsible for MHC I downregulation during influenza A virus infection. They also suggest that PA-X may hinder presentation of viral antigenic peptides to cytotoxic T cells, and thus help shelter the virus from the adaptive immune system.

## Discussion

In this study, we report that influenza A virus has evolved to use the same immunomodulatory protein, PA-X, to actively interfere with both the innate and adaptive arms of the immune responses. Using a combination of a primary human culture model that mimics the complexity of the human airway epithelium and scRNAseq, we carried a comprehensive analysis of different epithelial cell populations during infection, providing a broad view of relevant physiological changes. On one hand, PA-X significantly decreased signaling and/or secretion of type I and III IFNs, as well as acute pro-inflammatory cytokines and proteins associated with lung fibrosis and injury (Figures 2 and 3). These effects point to a role for PA-X in dampening initial inflammatory and activation signals in the context of innate immunity. On the other hand, PA-X disrupts expression of antigen processing and presentation genes and surface trafficking of MHC I, which can hinder presentation of new peptide antigens during infection and limit effector functions of CD8+ T cells. These dual activities likely explain the evolutionary pressure to retain PA-X through evolution.

The finding that PA-X reshapes cytokine secretion from the human airway epithelium during infection (Figure 3B) is in keeping previous results in mice. PA-X reduced pro-inflammatory cytokines IL-6, IL-1β, and G-CSF in human ALI culture (Figure 3B) and IL-6 and IL-1β in bronchoalveolar lavage of infected mice^29,32,37^. These cytokines are important for combatting influenza A viral infections, as patients with single nucleotide polymorphisms in IL-6 and IL-1β are more vulnerable to pandemic H1N1^77,78^. However, high levels of IL-6 and G-CSF are also predictors of influenza-associated pneumonia^113^. Pro-inflammatory cytokines must be carefully balanced during the infection response for this reason, and higher levels of these cytokines could explain the increased pathogenesis of PA-X deficient viruses in some studies^25,29–31,39^. PA-X regulation of type I and III IFN transcripts has also been reported in previous studies. However, at least in mice, previous studies have suggested there were no changes in interferon secretion (for example, Rigby et al.^36^). Instead, we show that in human epithelial cells, PA-X activity significantly decreases IFN-λ secretion. This is important because IFN-λ orchestrates the antiviral response in the airways^114^ and is induced by influenza A virus *in vivo*^68^ and in nasal epithelial cells^69^. Differences in IFN-λ secretion may have downstream effects on the adaptive immune response, as IFN-λ signaling is required for dendritic cell migration to draining lymph nodes and generation of protective memory immune responses to influenza A virus *in vivo*^115^. Reduction in IFN signaling could contribute to previously reported PA-X decrease in dendritic cell maturation and migration to draining lymph nodes *in vivo*^37,38^. More unexpected was the PA-X regulation of FGF basic and IGFBP-3. High levels of these factors are correlated with acute lung injury, acute respiratory distress, and pulmonary fibrosis^86–88^. As IGFBP-3 mediates extracellular matrix deposition^87^ and IL-6 contributes to fibrosis^116–118^, their regulation indicates that PA-X activity may have an impact on post-infection fibrosis, which would represent yet another facet of PA-X’s influence on virus-host interactions. This idea will need to be further tested with *in vivo* studies or complex human airway-fibroblast co-culture models^119^.

We were surprised to find that one of the strongest transcriptional signatures of PA-X activity, particularly in the infected cells, was not antiviral signaling but antigen presentation. To note, this was one of the pathways that Jagger et al. reported as reduced by PA-X in a mouse infection model^25^. MHC I antigen presentation allows the adaptive immune system, particularly CD8+ cytotoxic T cells, to detect cells infected with intracellular pathogens, and therefore it is an important obstacle for viruses to overcome. Our results reveal that antigen presentation mRNA downregulation by PA-X leads to reduced protein surface levels and delayed trafficking of MHC I. Downregulation of antigen presentation genes likely decreases MHC I trafficking both by directly lowering MHC I protein production and by reducing the efficiency of peptide loading, which can result in “empty” and/or unfolded MHC I complexes^105^. Indeed, PA-X cleaves the mRNAs for proteins like heat-shock protein 90 (encoded by *HSP90AA1*), which is needed to chaperone peptide antigens, and calreticulin (encoded by the gene *CALR*), which is needed to stabilize MHC I prior to peptide loading (Fig S7C-E). Our findings thus provide a mechanistic link between PA-X and disruption of MHC I protein expression and trafficking. Many viruses decrease surface MHC I during infection, including HIV^120–123^, adenoviruses^94,95^, herpesviruses^92,93^, papillomaviruses^96,97^, and coronaviruses^124,125^, some by reducing mRNA levels of MHC I complexes. Koutsakos et al. reported that influenza A viruses decrease surface and intracellular MHC I levels in infected A549 cells but did not determine the mechanism of MHC I decrease^20^. We now identify PA-X as the factor responsible for reduction of MHC I surface and intracellular levels by influenza A virus (Figure 4A-G). Through a reduction in antigen presentation, PA-X could help reduce cell targeting by antigen-specific CD8+ T cell response, allowing the virus to continue its spread. Importantly, the timing of MHC I changes in our experiments are consistent with a functional impact on peptide presentation to antigen-specific T cells. Presentation of influenza A virus antigen varies considerably between strains and cell types, as it mirrors the kinetics of influenza A virus protein production during replication^109^. This relationship was demonstrated by a classical study by Wu et al., which reported detection of MHC I bound influenza A virus antigens from mouse dendritic cells infected with the mouse adapted A/Puerto Rico/8/1934 H1N1 (PR8)^109^. In this study, peptides were detected as early as 3.5 hours post infection, reaching a peak by 6.5 hours, as were the proteins that those peptides came from^109^. However, in Calu-3 cells, Hamza et al. found that PR8 peptides accumulated later, in some cases peaking as late as 24 hours post infection^112^. Importantly the same study found that peptides produced during Calu-3 infection with a recent H3N2 (A/Fukui/20/2004) peaked as late as 24 hours post infection^112^. Similarly, we detected a reduction in MHC I trafficking to the cell surface of Calu-3 cells at 18-24 hours post infection (Figure 4F-G, Figure S8F-H). This comparison indicates that we captured an important window for presentation of peptide antigen during infection in Calu-3 cells. it is also important to note that our results on MHC I downregulation were consistent across 4 donors of different sexes, making it likely to occur in the human population during seasonal influenza infections (Figure 4D). In both our study and in Koutsakos et al.^20^ surface MHC I levels were reduced by 40% reduction, rather than completely eliminated. This partial reduction may have evolved as a compromise between avoiding detection and risking elimination of the infected cells by natural killer cells due to insufficient levels of surface MHC I. Suppression and/or delay of MHC I trafficking to the cell surface by PA-X could have broad implications for the success of cell-mediated immune responses following vaccination or during repeat infections. Influenza A virus is adept at causing seasonal illness despite pre-existing host immunity. Antigenic drift, i.e. the accumulation of mutations in viral surface proteins, is considered the predominant strategy of adaptive immune evasion by influenza A virus. However, antigenic drift does not account for significant reinfections with the same viral strain in human challenge studies^19^. Contrary to the more passive strategy of antigenic drift, here we reveal influenza A PA-X gene regulation as an active viral strategy for evasion of antigen-specific cell-mediated immunity via interference with MHC I. It is also intriguing to speculate that PA-X may have specifically evolved for this function in the context of pre-existing immunity to influenza A virus. Influenza A viral infection likely occurs predominantly in animals that have already encountered the virus, making evasion of adaptive immunity particularly important for the virus. A predominant role of PA-X in the context of subsequent infections would explain why the data in the literature on PA-X effects during infection in naïve animals are varied. Nonetheless, while our results point to an important new function for PA-X, further analysis of PA-X disruption of MHC I expression is needed to define its functional consequences on antigen presentation and CD8+ T cells. The functional effects of PA-X disruption of MHC I trafficking needs to be tested *in vivo*, to understand how this disruption affects T cell clearance of infected cells during primary infection and in immune hosts.

Beyond regulation of cytokine signaling and MHC I, our scRNAseq results point to additional PA-X effects on cell physiology and immune responses to be explored in future studies. PA-X may have specific effects on ciliated cells during infection, as PA-X activity promotes expression of genes involved in cilium assembly, yet decreases expression of nitric oxide production and cilia motility genes (Table 1B). Also, while we focused on downregulation of gene expression by PA-X, we also found that in a few cell types there were significantly enriched upregulated gene sets. In particular, in basal cells (clusters 0, 3) at 3 DPI, PA-X activity enriched expression of genes related to wound healing and development. This upregulation suggests that PA-X activity fosters the epithelial repair process, perhaps by decreasing inflammation, as excess or prolonged inflammation can delay wound healing^25,29,30,32,126^. Future studies could test whether PA-X has a protective effect on tissue repair and wound healing *in vivo.* Additionally, our dataset could be mined to analyze cell type-specific responses during influenza A virus infection in general. To conclude, here we have provided a roadmap for the study of PA-X in human airway epithelium and provided evidence for the importance of this poorly studied viral protein. Furthermore, our findings point to a new activity for PA-X in disruption of MHC I antigen presentation from infected cells, which we propose is the chief benefit of PA-X, allowing the virus to escape detection even in hosts with prior exposure.

### Limitations

Although we used a model relevant to human infections, our system only mimics the epithelial tissue, with no immune cells. While this study provides foundational knowledge that opens new avenues of investigation into a poorly understood but highly conserved influenza immunomodulator, many questions remain. In particular, future studies are needed to explore how the effect of PA-X on cytokines and MHC I impacts inflammation, antigen presentation of influenza A peptides, antigen-specific CD8+ T cell response and disease severity. *In vivo* influenza infections and/or more complex ALI co-culture models with different immune and non-immune cell populations will be necessary to define how the changes in cytokine signaling and MHC I expression affect the course of infection and disease.

## Materials and Methods

### Cell culture

Human bronchial epithelial (HBE) cells from 5 individual donors were used in our experiments. HBEs were procured from Lonza (Lonza Bioscience, CC-2540S) and a gift from J. Gern (University of Wisconsin-Madison, Madison, WI, USA)^127^. All primary human cells are de-identified, and therefore Institutional Review Board exempt. To generate ALI cultures, HBE cells were expanded using Pneumacult-Ex Plus media (STEMCELL Technologies) and seeded on collagen-coated 0.4 μm pore transwell inserts (Corning). Once cells reached confluency 2-3 days post seeding, the cultures were “air lifted” – i.e. media was removed from the apical chamber, exposing the cells to air. The basal chamber was filled with PneumaCult-ALI Medium (STEMCELL Technologies). Cells differentiated into a pseudostratified epithelium approximately 21 days post air lift. Cells were maintained in ALI culture at 37°C in 5% CO_2_ atmosphere. ALI cultures were used in experiments 3 weeks or more following air lift. Cultures were used for single-cell RNAseq 3-5 weeks post air lift.

Madin-Darby canine kidney (MDCK, ATCC cat# CCL-34) and Human embryonic kidney 293T (HEK293T, ATCC cat# CRL-3216) cells were used for virus production and titering. A previously described derivative of HEK293T with inducible knockdown of the human exoribonuclease Xrn1 (HEK293T ishXrn1)^128^ was used for the 5’ RACE experiments. MDCK, HEK293T, and HEK293T ishXrn1 cells were grown in Dulbecco’s modified Eagle’s medium (DMEM) high glucose (Gibco, cat# 11965118) (DMEM, Gibco) supplemented with fetal bovine serum (FBS, Hyclone) at 37°C in 5% CO_2_ atmosphere. Lung adenocarcinoma Calu-3 cells (ATCC, cat# HTB-55) were used for acid strip time course experiments. They were grown in 1X Minimum Essential Media (MEM, Gibco cat #11095080) supplemented with 10% FBS (Hyclone), 1X, GlutaMAX (Gibco, cat# 35050061), 1X Non-essential Amino Acid Solution (Gibco, cat# 11140050), 1% Pen/Strep (Gibco, cat# 15140122), and 1 mM Sodium Pyruvate (Gibco, cat# 11360070) at 37°C in 5% CO_2_ atmosphere.

| Donor | Source/Lot Number | Sex | Age | Race/ethnicity | Health conditions |
| --- | --- | --- | --- | --- | --- |
| 1 | Lonza/21TL244297 | M | 21 | Caucasian | None noted |
| 2 | Lonza/21TL347555 | M | 55 | Caucasian | None noted |
| 3 | Lonza/23TL023428 | M | 32 | Hispanic | None noted |
| 4 | Gern Lab | F | 62 | Caucasian | Subglottic stenosis |
| 5 | Gern Lab | F | 38 | Caucasian | Mild asthma |

### Plasmids and generation of viral stocks

Wild-type influenza A virus A/Perth/16/2009 H3N2 (Perth) and the mutant PA-X deficient virus (ΔX) were generated using the 8-plasmid reverse genetic system^129^ as previously described^45^. The rescue plasmids encoding the 8 Perth segments (pHW Perth PB2 to pHW Perth NS) were gifts from Dr. S. Lakdawala (Emory University School of Medicine, Atlanta, GA, USA^130^). The Perth pHW-PA(ΔX) plasmid was previously generated from pHW Perth PA by introducing mutations that reduce frameshifting events and add a premature stop codon in the PA-X reading frame, but that are silent in the PA reading frame^45,65^. Viral stocks were produced in MDCK cells and infectious titers were determined by plaque assays in MDCK cells using 1.2% Avicel (FMC BioPolymer) overlays^131^. For virus titering, confluent MDCK cells were infected with low volumes of tenfold serially diluted virus stocks in triplicate for 1 hour. Cells were then washed twice with DPBS before the addition of overlay media [1.2% Avicel, 2X MEM (Gibco), 0.5% w/v low-endotoxin bovine serum albumin (BSA, Sigma-Aldrich, cat # A2058) in DMEM (Gibco), and 1 ug/ml TPCK-treated trypsin (Sigma-Aldrich)] and incubated for 4 days at 37°C in 5% CO_2_ atmosphere. After 4 days, cells were fixed with 4% paraformaldehyde and stained with crystal violet (Sigma-Aldrich) to observe plaques.

### Viral infections

Influenza A virus infections of ALI cultures were performed in infection media (0.5% low-endotoxin BSA in high glucose DMEM). Briefly, differentiated ALI cultures were washed with Dulbecco’s phosphate buffered saline (DPBS, Gibco) for 20 minutes at 37°C in 5% CO_2_ atmosphere to remove excess mucus. ALI cultures were then treated with infection media alone (mock infection) or containing WT Perth or Perth PA(ΔX) at a multiplicity of infection (MOI) of 0.1 and incubated for 1 hour at 37°C in 5% CO_2_ atmosphere. The inoculum was then washed off. Every 24 hours, the apical side of each ALI culture was washed with 0.5% low-endotoxin BSA in DMEM for 20 minutes at 37°C in 5% CO_2_ atmosphere to collect virus. Media from the basal chamber was also collected and replaced at 24-hour intervals post infection to test cytokine secretion.

### Confocal microscopy

Infected or mock-infected ALI cultures were fixed in transwells with 4% paraformaldehyde for 30 minutes, then stored in DPBS (Gibco) at 4°C. ALI culture transwell inserts were permeabilized and blocked by adding 0.1% Triton X100 and 1% BSA (Thermo Fisher Scientific) to the apical chamber and incubating on a rocker at room temperature for 30 minutes. Each transwell insert was cut into multiple pieces (3-4) to allow for multiple antibody stains from each ALI culture. Transwell pieces were placed in a 96-well plate for staining. Primary antibody solutions were prepared in 0.1% Triton X100 and 1% BSA, and each transwell section was stained in 100 μl of primary antibody solution for 1 hour on a rocker at room temperature. Fixed ALIs were washed 3 times with DPBS (Gibco) and then stained with secondary antibodies at a 1:500 dilution. Nuclei were stained with 1:10,000 dilution of Hoechst 3342 Fluorescent Stain (Thermo Fisher Scientific) for 30 minutes in the dark. Cells were washed again with DPBS (Gibco) and mounted on slides using ProLong Gold Antifade Mountant (Invitrogen). Images and z-stacks were taken with a Leica Stellaris Confocal Microscope, and sagittal views were computed using ImageJ^132^. The following antibodies were used:

| Target | Stains for | Clone | Cat no. | Company | Produced in | Dilution | Fluor |
| --- | --- | --- | --- | --- | --- | --- | --- |
| ZO-1 | Tight junctions | ZO1-1A12 | 33-9100 | Thermo Fisher | mouse | 1:50 | AF488 |
| Acet-TUB1a | Cilia | 6-11B-1 | MABT868 | Millipore Sigma | mouse | 1:100 | AF488 |
| MUC5AC | Mucin 5AC | 45M1 | ab3649 | abcam | mouse | 1:100 | AF594 |
| Influenza A virus nucleoprotein (NP) | Influenza A virus | HB-65 | BE0159 | Bioxcell | mouse | 1:100 | AF680 |

Secondary Alexa-Fluor conjugated antibodies were purchased from Thermo Fisher Scientific. NP was pre-conjugated using Antibody Labeling Kits for 1 mg (Thermo Fisher Scientific).

### Single cell RNA sequencing (ScRNAseq)

#### Cell collection, library generation, and sequencing

ALI cultures from donors 1 and 2 were collected 1 and 3 days post infection for scRNAseq. Cells were dissociated with trypsin (Lonza) in the presence of DNase I (Thermo Fisher Scientific) and Protector RNase Inhibitor (Millipore Sigma) for 30 minutes. Dead cells were depleted using EasySep Dead Cell Removal Annexin V Kit (STEMCELL Technologies), and a viable single cell suspension was fixed for library prep using the Evercode Cell Fixation Kit (Parse Biosciences) according to the manufacturer’s protocol. Library prep was performed at the University of Wisconsin-Madison Gene Expression Center using WT Evercode v3 (Parse Biosciences) according to the manufacturer’s protocol. During library preparation, cells were split to generate 8 sublibraries, which were sequenced on the NovaSeq X Plus 10B 2×150 flow cell at the University of Wisconsin-Madison Gene Expression Center. Approximately 6 billion reads (1.25B +/- 10% per lane, 5 lanes) were collected. About 69,000 cells were detected, with a depth of approximately 20,000 reads/cell.

### Alignment, QC, Data processing

Raw Parse Evercode WT sequencing data was processed per sublibrary by the Parse Evercode pipeline (split-pipe version 1.3.0), aligning to the human genome (GRCh38) primary assembly with Ensembl v112 annotations and influenza A virus Perth H3N2 genome (Genbank Accessions: KJ609203-KJ609210). Data was downloaded from Trailmaker for further processing in the R program Seurat (version 5.1.0), as described by Parse Biosciences. Briefly, a Seurat object was created by reading in the Trailmaker-generated “DGE_filtered” data, which contains gene lists, cell metadata, and a count matrix. Cells were then filtered for quality by excluding cells with >25% mitochondrial RNA, >12k RNA features, and >20k RNA reads per cell, which likely includes dead cells.

To minimize variation due to donors and processing, the Seurat object was integrated using the Seurat functions FindIntegrationAnchors() and IntegrateData(). Unbiased clustering analysis was performed using the Seurat package. Cell type was assigned with ScType^52^ and then additionally curated using additional markers from the literature. Pseudo time analysis was performed using Monocle 3^133–135^, as described by the Trapnell lab. Scripts used for analysis can be found on Github.

### Differentially Expressed Genes (DEGs) and Gene Set Enrichment Analysis

The Seurat program FindAllMarkers was used to find DEGs between different conditions. DEGs were defined as genes with |log2FC| ≥ 0.25 and p-adjusted <0.05. These DEGs were then used to determine enriched gene sets with the clusterProfiler command enrichGO( ) for gene ontology over representation^58,136–138^. We then reduced redundancy using the simplify( ) method in clusterProfiler. For all gene enrichment, the human genome database org.Hs.eg.db was used. Gene enrichment results were visualized using the DOSE package^59^. Scripts used for analysis can be found on Github.

### Flow cytometry analysis

#### Epithelial cell subtyping

ALI cultures grown in 12-mm transwell inserts were collected at indicated times post infection. ALI cultures were trypsinized for 20-30 minutes at 37°C, then the reaction was neutralized with Trypsin Neutralization Solution (Lonza). Cells were stained with fixable viability Violet (Thermo Fisher Scientific) for 30 minutes in the dark on ice. Cells were then fixed with 4% paraformaldehyde, then resuspended in flow buffer: 1X PBS, no calcium or magnesium (Fisher Bioreagents), 1mM EDTA (Fisher Bioreagents), 25 mM HEPES (Cytiva Hyclone), 2% FBS (Cytiva). Flow buffer was also used in all subsequent wash and staining steps, as FBS decreases non-specific staining. Cells were permeabilized with 0.5% Tween 20 (Thermo Fisher Scientific) and 1% w/v BSA (Thermo Fisher Scientific) in 1X PBS (Fisher Bioreagents) for 20 minutes and stained with appropriate antibodies. Additional wells were harvested for single color and full minus one (FMO) controls for each fluorophore. Cells were analyzed on the Cytek Aurora (Cytek) at the University of Wisconsin-Madison flow cytometry core. To deconvolute overlapping emission spectra of different fluorophores, unmixing was performed on the SpectroFlo software (Cytek). Unmixing was performed using cell controls, as we found unmixing with bead controls was ineffective likely due to background fluorescence of the cells. The following antibodies were used:

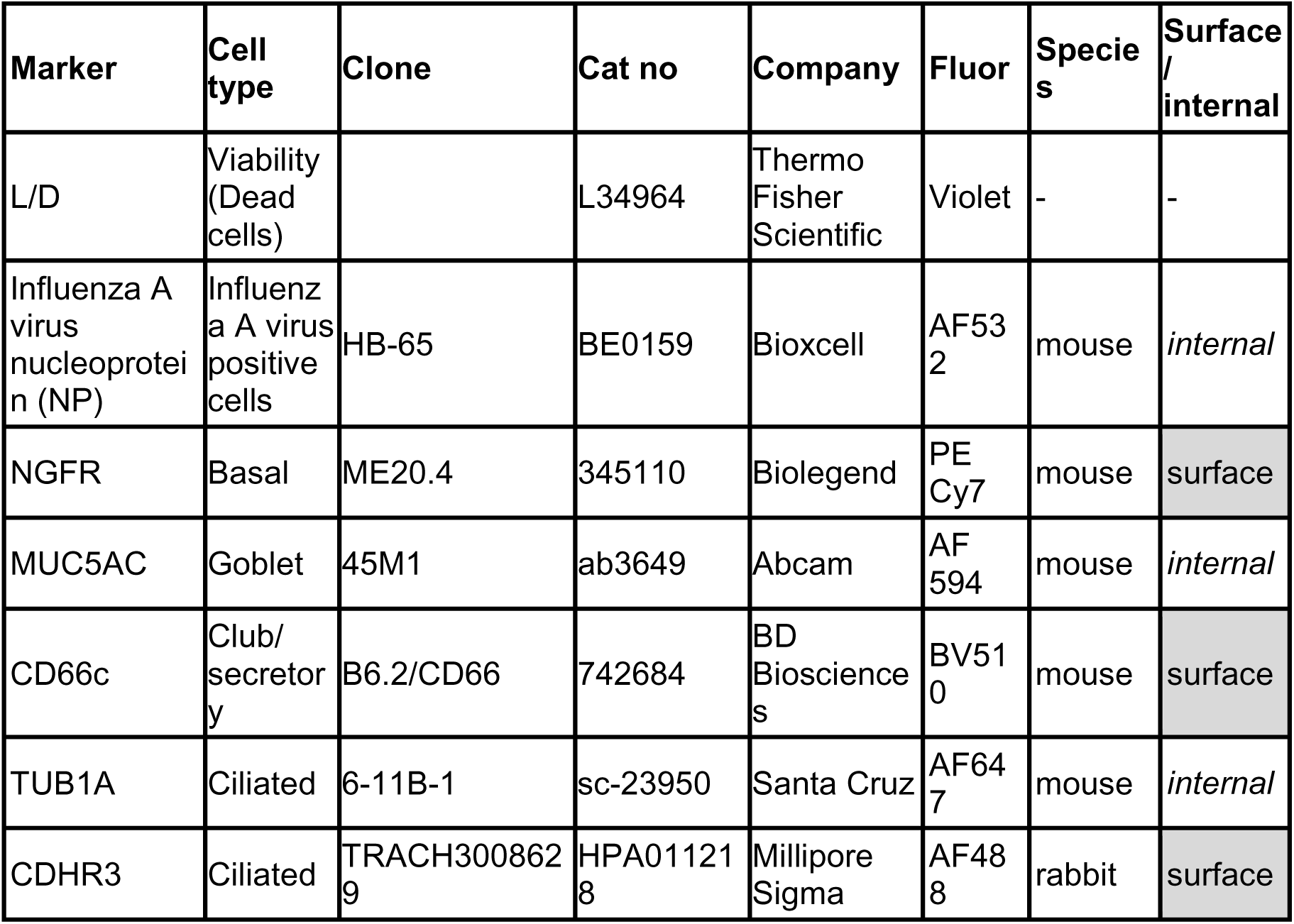

Antibodies against NGFR, CD66C, and TUB1A were purchased pre-conjugated to their fluors. NP and MUC5AC were conjugated to Alexa Fluor 532 and Alexa Fluor 594, respectively, in the laboratory using Antibody Labeling Kits for 1 mg (Thermo Fisher Scientific). Secondary antibody anti-Rabbit Alexa Fluor 488 (Thermo Fisher Scientific) was used to detect CDHR3 staining.

### Analysis of MHC I levels in ALI cultures

ALI cultures grown in 6.5 mm transwell inserts were collected each day post infection by trypsinizing for 20-30 minutes at 37°C, neutralizing the reaction with Trypsin Neutralization Solution (Lonza) and staining cells with Live Dead Violet (Thermo Fisher Scientific) for 30 minutes in the dark on ice. Cells were fixed with 4% paraformaldehyde and then resuspended in flow buffer (described above). All following wash and staining steps were performed in flow buffer. Cells were stained for surface MHC I using HLA-ABC W6/32 (eBiosciences, 14-9983-82) conjugated to Alexa Fluor 488 or FITC, depending on the experiment, for 45 minutes at room temperature on a rocker in the dark. After MHC I surface stain was washed off, cells were permeabilized with 0.5% Tween 20 (Thermo Scientific) for 20 minutes. Cells were washed and then stained with MHC I HLA-ABC W6/32 pre-conjugated to Alexa Fluor 594 and Influenza A Virus NP antibody (clone HB-65; Bioxcell BE0159) pre-conjugated with Alexa Fluor 680. Conjugation was performed using Antibody Labeling Kits for 1 mg (Thermo Fisher Scientific). Cells were analyzed on the Attune NxT (Thermo Fisher Scientific) at the University of Wisconsin-Madison flow cytometry core.

### Analysis of MHC I levels during the acid strip time course

Calu-3 cells were seeded in a 6-well plate and grown to 80-90% confluence. Cells were either mock-infected or infected with WT or ΔX Perth at MOI 0.1. After incubation for 1 hour at 37°C in 5% CO_2_ atmosphere, the inoculum was removed and replaced with Calu-3 infection media (Calu-3 media, described in the Cell Culture section, without FBS, and supplemented with 0.5% w/v low-endotoxin BSA). 18 hours post infection, cells were trypsinized, then neutralized with Calu-3 media. An initial pre-strip sample was collected at this time and placed on ice. Cells from each condition were spun down at 300 x *g*. Cell pellets (with approximately 1 million cells) were acid stripped using 200 μl of citric acid-Na_2_HPO_4_ buffer for 2 minutes at 4C. Citric acid-Na_2_HPO_4_ buffer was prepared with equal volumes of 0.263 M citric acid and 0.123 M Na_2_HPO_4_ with 1% BSA (w/v), which was then titrated to a pH of 3.0^110^. The acid was neutralized with 14.8 ml of Calu-3 media. Cells were spun down at 300 x *g* and resuspended in Calu-3 media. The 0-hour post strip time point was collected on ice. The remaining cells were plated in a 12-well plate and collected at 2, 4, and 6 hours post acid strip. After the collection of each time point, cells were collected on ice and then stained with Live Dead Violet (Thermo Fisher Scientific) for 30 minutes in the dark on ice. Cells were fixed with 4% paraformaldehyde and then resuspended in flow buffer (described above). Cells were stained for surface MHC I using HLA-ABC W6/32 conjugated to FITC (eBiosciences, 14-9983-82) for 45 minutes at room temperature on a rocker in the dark. After the MHC I surface stain was washed off, cells were permeabilized with 0.5% Tween 20 (Thermo Scientific) for 20 minutes, washed, and stained with Influenza A Virus NP antibody (clone HB-65; Bioxcell BE0159) pre-conjugated with Alexa Fluor 680. Conjugation was performed using the Antibody Labeling Kit for 1 mg (Thermo Fisher Scientific). Cells were analyzed on the Attune NxT (Thermo Fisher Scientific) at the University of Wisconsin-Madison flow cytometry core.

### Measurement of IFN secretion using reporter cells and enzyme linked immunosorbent assay (ELISA)

HEK-Blue™ IFN-λ Cells (InvivoGen) were used to quantify type III interferon in media from the basal chamber of ALI culture wells. HEK-Blue™ IFN-λ Cells were cultured according to the InvivoGen’s protocol. Briefly, cells were expanded for two passages in DMEM with 2 mM L-glutamine (Gibco) with 10% FBS (Cytiva). Cells were then grown under antibiotic selection in media supplemented with 10 µg/ml of Blasticidin (InvivoGen), 1 µg/ml of Puromycin (InvivoGen) and 100 µg/ml of Zeocin (InvivoGen).

Following antibiotic selection, HEK-Blue™ IFN-λ Cells were detached with DPBS and resuspended in growth medium without selection antibiotics, then added to a flat-bottomed 96-well plate. Conditioned media from the basal chamber of ALI cultures was then added to the flat-bottomed 96-well plate. To ensure that the cells were specific for IFN-λ, human interferon alpha 2b (InvivoGen) was used as a negative control, and a standard was generated using Human IL-28A (IFN-lambda 2, PeproTech) as a positive control. All samples and controls were tested in triplicate. Samples were incubated with HEK-Blue™ IFN-λ Cells overnight at 37°C in 5% CO_2_ atmosphere. The following day, HEK-Blue™ IFN-λ Cell supernatant was incubated with QUANTI-Blue™ solution (InvivoGen) at 37°C in 5% CO_2_ atmosphere for 30 minutes to 3 hours, then plates were read on the Cytation 5 plate reader (BioTek) at 620 and 655 nm. Protein levels were calculated based on Human IL-28A standard curve using non-linear least squares calculated on R. This code is available on GitHub (fittingdata.R). IFN-β secretion in conditioned media from the basal chamber of ALI cultures was measured using Human IFN-beta bioluminescent ELISA kit 2.0 (Invivogen) according to manufacturer instructions.

### Measurement of cytokine secretion using Luminex

Protein levels were quantified on the Luminex MAGPIX Analyzer using a custom panel including color coded magnetic microspheres with capture antibodies, as well as biotinylated detection antibodies for each analyte (R&D Systems, LXSAHM-18). To select panel targets, we took PA-X dependent DEGs from our dataset and entered them into the web application CytoSig. CytoSig predicts cytokines that may be altered by analyzing transcriptional signatures in sequencing data^84^. We narrowed down this initial list by excluding targets that were not expressed in our scRNAseq dataset and by selecting ones for which there were commercially available Luminex kits. Samples of conditioned basal media were vortexed and spun down for 10 minutes at 1000 x *g* prior to performing Luminex assay according to manufacturer protocol. Donors 2-5 samples and standards were tested in technical duplicate, while Donor 1 samples which were tested in single replicates due to limited reagent and space on the plate. The assay was read on the Luminex MAGPIX Analyzer and protein concentrations were calculated on the xPONENT® program (UW Madison Vision Core).

### 5’ rapid amplification of cDNA ends (RACE)

Direct cleavage of RNAs was measured using 5’ RACE, as described previously^65,139^. Briefly, HEK293T ishXRN1 cells were treated with 1 μg/mL doxycycline for 3-4 days to induce the XRN1 shRNA, knock-down the protein, and stabilize RNA fragments generated by PA-X. They were then transfected with Perth H3N2 PA-X (pCR3.1-H3N2 Perth PA-X-myc^65^), Perth H3N2 PA-X D108A (pCR3.1-H3N2 Perth PA-X-D108A-myc, generated by Gibson cloning), or empty vector, together with pd2eGFP-N1 (to monitor transfection efficiency) and the pCMV-luciferase+intron-YKT6-99bp cleavage reporter^65^. RNA was extracted from cells using the Quick-RNA miniprep kit (Zymo Research) following the manufacturer’s protocol and treated with Turbo Dnase (Life Technologies). After phenol/chloroform extraction from the DNase reaction, the RACE adapter (sequences below) was ligated to RNA using T4 RNA ligase (Ambion) at 25°C for 2 hrs and reverse transcribed using MMLV RT (Invitrogen). Fragments of interest were amplified by Taq DNA polymerase (New England Biolabs) using forward primers annealing to the RACE adapter and reverse primers annealing to the transfected reporter construct (sequences below^65,128^). PCR products were separated on a 2% agarose gel containing HydraGreen safe DNA dye (ACTGene) and visualized using the iBright FL1000 imager system and LI-COR Odyssey XF. DNA was extracted from gel bands at expected sizes and Sanger sequenced to confirm their identities.

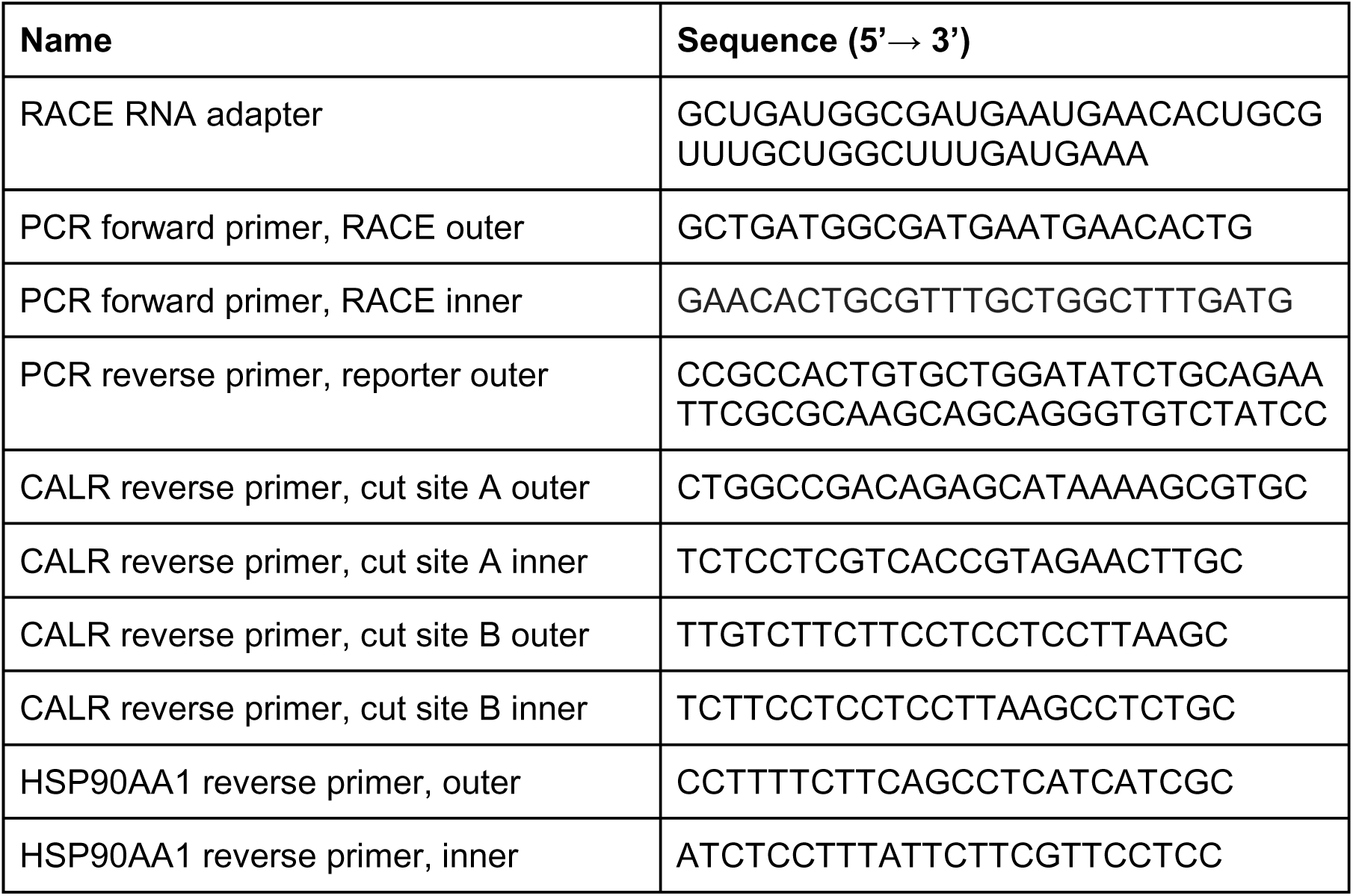

### Statistical analysis

Unless otherwise noted, data represent three or more independent experiments using cells from different donors. Bar plots with error bars represent mean ± standard deviation. For box and whiskers, boxes extend from the first quartile to the third quartile, with horizontal lines denoting the median value and whiskers extending to minimum and maximum values. Floating bar graphs have boxes extending from minimum to maximum values with horizontal line denoting the median. Statistical analyses were performed using GraphPad Prism (version 10). One-way or two-way analysis of variance (ANOVA) with Dunnett’s multiple comparison test were used for multiple comparisons. To account for high levels of donor variability in cytokine secretion during WT infection, ratio paired t tests were used to test differences between WT and ΔX infection. For acid strip experiments, the fold-change in levels compared to the 0-hours post acid strip sample was plotted. Linear regression and statistics were performed using GraphPad Prism. Slopes were compared on GraphPad Prism, using a technique equivalent to one form of ANCOVA.

## Supporting information

All supplemental materials

## Data availability

Raw and processed data is available on NCBI GEO: GSE319472. Scripts used for analysis are available on GitHub (https://github.com/mgaglia81/HBE_ALI_scRNAseq). Primary data for the final version of the manuscript will be made available on Figshare upon publication (private reviewer link: https://figshare.com/s/8e7444b3fd99c8222550).

## Acknowledgments

We thank members of the Gaglia laboratory for suggestions and feedback on the project and the manuscript. We thank M. Hing and J. Feng for feedback and technical support on figures. We thank the University of Wisconsin Carbone Cancer Center Flow Cytometry Laboratory, (supported by P30 CA014520 and Cytek Aurora Spectral Cytometer, “Pinky” Grant #: 1S10OD025225-01), for use of its facilities and services. We thank the UW Madison vision core, (supported by P30 EY016665), for training on and usage of the Luminex MAGPIX. We thank K. Majumder for use of the confocal microscope, S. Lakdawala for constructs, and J. Gern for sharing donor cells. This work was supported by Emory CEIRR 75N93021C00017 Option 16E (to MMG), NIH R01 AI137358 (to MMG), and the University of Wisconsin-Madison Office of the Vice Chancellor for Research and Graduate Education with funding from the Wisconsin Alumni Research Foundation.

## Author Contributions

A.C.S. and M.M.G. designed research; A.C.S. and C.A.T. performed research; A.C.S. analyzed data; and A.C.S. and M.M.G. wrote the paper.

## Competing Interest Statement

The authors declare no competing interests.

---

**Supplemental Table 3. PA-X does not alter infection parameters at the single cell level.** ALI cultures were infected and samples collected as described in Figure 1D. The scRNAseq data were analyzed to identify infected cells (n = 2, Donors 1 and 2). A) Integrated UMAP of mock, Perth H3N2 WT and Perth H3N2 ΔX infected ALI cultures colored by the percent of reads that mapped to influenza A viral genes in each cell. B) Bar graph showing the percentage of infected cells that belong to each epithelial cell type vs. the composition of mock-infected cultures. Cells were deemed infected if at least 1% reads mapped to the influenza transcriptome. The two donors were aggregated for the analysis in all panels. The two timepoints (1 and 3 DPI) were aggregated for panel A.

**Figure S1. Airway epithelial cell subsets can be identified by transcriptional markers.** A-B) ALI cultures were infected and samples collected as described in Figure 1D. The scRNAseq data were analyzed to identify groups of cells with similar expression patterns and to assign cell type (n = 2, Donors 1 and 2). A) (Top left, same as Figure 1E) Integrated UMAP diagram of all samples, showing the 15 clusters unbiasedly identified by the program Seurat. (Top right, same as Figure 1F) Integrated UMAP diagram of all samples displaying epithelial cell types assigned with ScType. The clusters mapping to each cell type are listed in parentheses. (Below) Feature plots show the expression levels of markers associated with different cell types and subsets. FOXN4 marks the ciliated cell cluster 14 specifically as deuterosomal ciliated cells. The expression of POUF2F3 and CHGA in cluster 13 shows that this cluster likely includes not only ionocytes but also tufts cells and pulmonary neuroendocrine cells (PNECs), two rare subtypes of epithelial cells. B) UMAP showing epithelial cell differentiation pathways with pseudotime analysis in all donors, timepoints, and conditions using Monocle3. Pseudotime begins at 0, representing the earliest point in differentiation, and increases to represent further differentiation from the starting point. As expected pseudotime 0 corresponds to basal cells, which are the stem cells of the airway. While clusters 2 and 11 were previously undetermined by ScType, pseudotime analysis helped us to determine that these clusters are likely basal cells that have begun to differentiate into other airway epithelial cell types. Black circles represent nodes where cell trajectory branches into several different outcomes. Numbers inside each node are for reference purposes only, and do not hold other meaning.

**Figure S2. Influenza A viral infection alters cell type composition in the airway epithelium in a PA-X-independent manner.** ALI cultures were infected and samples collected as described in Figure 1D. scRNAseq data from Perth H3N2-infected ALI cultures were analyzed to identify groups of cells with similar expression patterns and to assign cell type (n = 2, Donors 1 and 2). A) Human bronchial epithelial cells were cultured at ALI for 3-4 weeks and stained for tight junctions (ZO-1, top), cilia (acetylated TUB1A, middle, sagittal section), and the mucus component and goblet cell marker mucin 5AC (MUC5AC, bottom) and nuclear stain (Hoescht, all) to confirm proper differentiation of the cells and formation of the pseudostratified epithelium. B) Immunofluorescence image of infected ALI culture, with staining of influenza A virus nucleoprotein (NP), cilia (acetylated TUB1A), and nuclear stain (Hoechst) at 2 days post infection. C) Cells were collected for flow cytometric validation of cell types at 1 day post infection (DPI). (Left) Gating strategy for epithelial cell types in ALI cultures after gating for live, single cells. We termed cells negative for ciliated, secretory, and basal markers “triple-negative” cells. They likely represent the cells in cluster 13 (Figure 1E). (Right) Stacked bar plot of average epithelial cell type composition from two different donors (Donors 4, 5). D) Alluvial plot showing the change in proportions of cells per cluster and cell type throughout infection with Perth H3N2 WT and ΔX. The two donors are plotted separately. The percentage of dead cells was calculated based on cell viability measurements using trypan blue carried out prior to dead cell depletion for scRNAseq library preparation.

**Figure S3. PA-X does not alter infection parameters at the single cell level.** ALI cultures were infected and samples collected as described in Figure 1D. The scRNAseq data were analyzed to identify infected cells (n = 2, Donors 1 and 2). A) Integrated UMAP of mock, Perth H3N2 WT and Perth H3N2 ΔX infected ALI cultures colored by the percent of reads that mapped to influenza A viral genes in each cell. B) Bar graph showing the percentage of infected cells that belong to each epithelial cell type vs. the composition of mock-infected cultures. Cells were deemed infected if at least 1% reads mapped to the influenza transcriptome. The two donors were aggregated for the analysis in all panels. The two timepoints (1 and 3 DPI) were aggregated for panel A.

**Figure S4. Influenza A virus infected cells arrest in G0/G1, while bystander cells do not.** Analysis of scRNAseq data from ALI cultures infected as shown in Figure 1D (n = 2, Donors 1 and 2). A-B) Bar graphs showing analysis of cell cycle phase using Seurat at 1 day post infection (DPI), shown according to A) cluster and assigned cell type or B) infection status. Infected cells = influenza A virus genes ≥1% of total reads; uninfected cells = influenza A virus genes < 1% of total reads.

**Figure S5. PA-X does not alter IFN-β secretion during infection.** ALI cultures were infected with Perth H3N2 WT or ΔX or mock infected at MOI 0.1. Basal media was collected every day after infection and IFN-β protein levels were tested using ELISA. A) IFN-β protein concentration in basal media. C) Log_2_ fold-change of IFN-β secreted by ΔX-infected ALIs vs. WT-infected (n = 3 experiments in donor 1).

**Figure S6. PA-X limits secretion of pro-inflammatory and lung injury-associated cytokines** A-B) Luminex data using basal conditioned media from ALI cultures infected as shown in Figure 1D (n = 5, 5 separate donors, Donors 1-5). B) Bar graphs showing concentrations of cytokines throughout infection time course. Comparisons between conditions are shown (analyzed using a two-way ANOVA, followed by Šídák’s multiple comparisons test. ns = p > 0.05, *= p ≤ 0.05, **= p ≤ 0.01, ***= p ≤ 0.001.) B) For targets that were outside the limit of detection (LOD) of the standard curve, median fluorescence intensity is plotted in bar graphs. Comparisons between conditions are shown (analyzed using a two-way ANOVA. ns = p > 0.05, *= p ≤ 0.05, **= p ≤ 0.01, ***= p ≤ 0.001.)

**Figure S7. PA-X directly cleaves antigen processing and presentation genes.** A) Schematic of the 5’ RACE workflow. 5’ P, 5’ phosphate. B) Table of cut sites detected by 5’ RACE-seq analysis in Gaucherand et al. 2023 in the significantly downregulated genes listed in Figure 4B. Arrows indicated the sites that were selected for experimental validation. C-E) XRN1 inducible knockdown HEK 293T cells were transfected with empty vector, WT Perth H3N2 PA-X, or catalytically inactive (D108A) Perth H3N2 PA-X for 24 hours before RNA extraction. 5’ RACE was then performed using primers downstream of cut sites identified in the 5’ RACE-seq in (C-D) CALR and (E) HSP90AA1. Arrows indicate the position of PCR fragments that correspond to the expected cut sites. The DNA bands were purified and sequenced to confirm their identities. Additional bands in control samples are derived from RNA fragments present in the cell but unrelated to PA-X activity.

**Figure S8. MHC I levels during infection.** A-C) ALI cultures were infected with Perth H3N2 WT or ΔX virus, or mock-infected at MOI 0.1. Fixed cells were stained with an anti-HLA-ABC antibody prior to permeabilization to detect the surface exposed portion of MHC-I. The cells were then permeabilized and stained for internal HLA-ABC, using the same antibody coupled with a different fluorophore. Cells were also stained for influenza A virus nucleoprotein (NP). IAV+ = influenza A virus NP-positive cells. A) Infected cells NP+ cells at 1 and 3 days post infection (DPI) as a percent of total live cells. B) Median fluorescence intensity of surface HLA-ABC. C) Median fluorescence intensity of intracellular HLA-ABC. D-E) ALI cultures were infected with Cal H1N1 WT or ΔX virus, or mock-infected at MOI 0.1. Cells were stained as described for A-C. D) HLA-ABC median fluorescence intensity at 2 DPI on NP+ infected cells from flow cytometry analysis plotted as fold-change relative to mock-infected ALI cultures. E) Infected cells NP+ cells at 1-3 DPI as a percent of total live cells. F-H) Calu-3 cells were mock-infected or infected with Perth H3N2 WT or ΔX at an MOI of 0.1 for 18 hours. Surface MHC I was stripped using a citric acid buffer, then cells were collected at multiple timepoints over the course of 6 hours. F) Median fluorescence intensity of MHC I on the surface of Calu-3 cells during this experiment. G) Percent of influenza NP+ cells. H) Log_2_ fold change of surface MHC I median fluorescence intensity over a 6-hour time course in influenza A virus NP- bystander cells in WT and ΔX-infected cultures. Linear regression was performed for each condition, and the p value was calculated to compare Perth H3N2 WT and ΔX NP- bystander cells. Dashed lines represent the line of best fit, and the shaded regions indicate the 95% confidence interval (see Figure 4F for equivalent plot for mock-infected cells).

## Notes

### Competing Interest Statement

The authors have declared no competing interest.

### Summary of Updates

Revised the title an the text to moderate the language on some of the conclusions in response to reviewers comments.

