## Supplementary material for "Dual effects of influenza A virus PA-X on suppression of cytokine signaling and MHC I pathway disruption in the human airway epithelium": All supplemental materials

**Supplemental Table 1. Donor information**

| Donor number | Sex | Age | Ethnicity | Health conditions |
|---|---|---|---|---|
| 1 | M | 21 | Caucasian | None noted |
| 2 | M | 55 | Caucasian | None noted |
| 3 | M | 32 | Hispanic | None noted |
| 4 | F | 62 | Caucasian | Subglottic stenosis |
| 5 | F | 38 | Caucasian | Mild asthma |

**Supplemental Table 2. Donors used in each experiment**

| Experiment | Panel | Donor used |
|---|---|---|
| Viral replication | Figure 1E | 1, 2, 3, 4, 5 |
| Barrier integrity | Figure 1F | 2, 3, 4 |
| Single cell RNAseq | Figure 2 A-B, Figure 3, Table 1, Figure 4 A-B, D, Figure 6 A-B | 1, 2 |
| Epithelial cell flow cytometry | Figure 2 C | 4, 5 |
| IFN III secretion | Figure 4 C | 1, 2, 3, 4, 5 |
| HLA-ABC flow cytometry | Figure 6 C, D | 2, 3, 4, 5 |

| Supplemental Table 3. Different subtypes of cells respond differently to influenza infection | | | |
|---|---|---|---|
| **Gene set enrichment analysis: Upregulated gene sets in WT infection versus mock** | | | |
| **Cell type** | **Cluster** | **1 DPI** | **3 DPI** |
| **Airway goblet cells** | **7** | NA | epithelial cell proliferation<br>response to interleukin-1<br>granulocyte migration |
| | **8** | NA | microtubule bundle formation<br>keratinization<br>cytokine activity |
| **Basal cells** | **0** | presynapse organization<br>apical plasma membrane | positive regulation of phosphorylation<br>granulocyte chemotaxis<br>humoral immune response |
| | **3** | axoneme<br>ciliary plasm<br>cytoplasmic microtubule | humoral immune response<br>neutrophil migration<br>regulation of wound healing |
| | **9** | NA | cell adhesion mediated by integrin<br>negative regulation of viral genome replication<br>response to lipopolysaccharide |
| **Basal cells (Ki67+)** | **12** | microtubule bundle formation<br>axoneme assembly<br>secretory granule membrane | response to lipopolysaccharide<br>leukocyte migration<br>humoral immune response |
| **Ciliated cells** | **5** | mitotic sister chromatid segregation<br>cellular response to interleukin-1<br>microtubule motor activity | myeloid leukocyte migration<br>response to lipopolysaccharide<br>response to interleukin-1 |
| | **14** | NA | NA |
| **Ciliated cells (infected)** | **10** | B cell differentiation<br>cell chemotaxis | leukocyte migration<br>cellular response to fibroblast growth factor stimulus<br>response to interleukin-1 |
| **Club cells** | **1** | monoatomic ion channel complex<br>histone H3Y41 kinase activity | wound healing<br>lung development<br>myeloid leukocyte migration |
| | **4** | NA | microtubule bundle formation<br>myeloid leukocyte migration<br>response to fibroblast growth factor |
| **Club cells (infected)** | **6** | microtubule-based movement<br>cellular response to lipopolysaccharide<br>cytokine activity | myeloid leukocyte migration<br>cell-cell junction assembly<br>neutrophil chemotaxis |
| **Ionocytes** | **13** | NA | microtubule-based movement<br>collagen-containing extracellular matrix<br>cytokine activity |
| **Basal-like cells** | **2** | mitotic spindle organization<br>chromosome segregation<br>microtubule motor activity | leukocyte migration<br>neutrophil migration<br>wound healing |
| | **11** | postsynaptic membrane | microtubule bundle formation<br>cilium or flagellum-dependent cell motility<br>multicellular organismal response to stress |

**Supplemental Table 3. PA-X does not alter infection parameters at the single cell level.** ALI cultures were infected and samples collected as described in Figure 1D. The scRNAseq data were analyzed to identify infected cells (n = 2, Donors 1 and 2). A) Integrated UMAP of mock, Perth H3N2 WT and Perth H3N2 ∆X infected ALI cultures colored by the percent of reads that mapped to influenza A viral genes in each cell. B) Bar graph showing the percentage of infected cells that belong to each epithelial cell type vs. the composition of mock-infected cultures. Cells were deemed infected if at least 1% reads mapped to the influenza transcriptome. The two donors were aggregated for the analysis in all panels. The two timepoints (1 and 3 DPI) were aggregated for panel A.

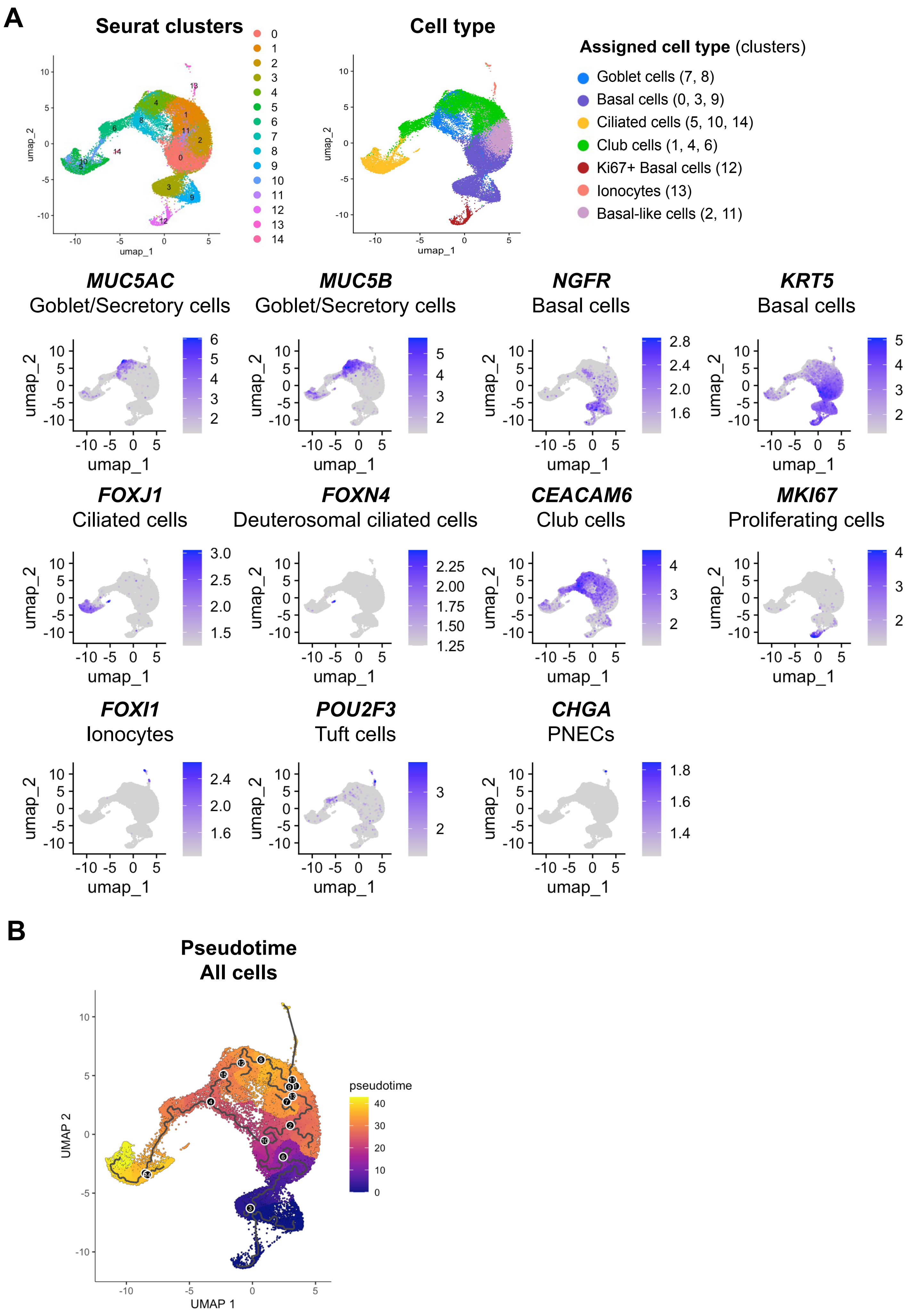
A
Seurat clusters
0 1 2 3 4 5 6 7 8 9 10 11 12 13 14
umap_1
umap_2
Cell type
Assigned cell type (clusters)
Goblet cells (7, 8)
Basal cells (0, 3, 9)
Ciliated cells (5, 10, 14)
Club cells (1, 4, 6)
Ki67+ Basal cells (12)
Ionocytes (13)
Basal-like cells (2, 11)
MUC5AC
Goblet/Secretory cells
MUC5B
Goblet/Secretory cells
NGFR
Basal cells
KRT5
Basal cells
FOXJ1
Ciliated cells
FOXN4
Deuterosomal ciliated cells
CEACAM6
Club cells
MKI67
Proliferating cells
FOXI1
Ionocytes
POU2F3
Tuft cells
CHGA
PNECs
B
Pseudotime
All cells
pseudotime
UMAP 1
UMAP 2

**Figure S1. Airway epithelial cell subsets can be identified by transcriptional markers.** A-B) ALI cultures were infected and samples collected as described in Figure 1D. The scRNAseq data were analyzed to identify groups of cells with similar expression patterns and to assign cell type (n = 2, Donors 1 and 2). A) (Top left, same as Figure 1E) Integrated UMAP diagram of all samples, showing the 15 clusters unbiasedly identified by the program Seurat. (Top right, same as Figure 1F) Integrated UMAP diagram of all samples displaying epithelial cell types assigned with ScType. The clusters mapping to each cell type are listed in parentheses. (Below) Feature plots show the expression levels of markers associated with different cell types and subsets. FOXN4 marks the ciliated cell cluster 14 specifically as deuterosomal ciliated cells. The expression of POUF2F3 and CHGA in cluster 13 shows that this cluster likely includes not only ionocytes but also tufts cells and pulmonary neuroendocrine cells (PNECs), two rare subtypes of epithelial cells. B) UMAP showing epithelial cell differentiation pathways with pseudotime analysis in all donors, timepoints, and conditions using Monocle3. Pseudotime begins at 0, representing the earliest point in differentiation, and increases to represent further differentiation from the starting point. As expected pseudotime 0 corresponds to basal cells, which are the stem cells of the airway. While clusters 2 and 11 were previously undetermined by ScType, pseudotime analysis helped us to determine that these clusters are likely basal cells that have begun to differentiate into other airway epithelial cell types. Black circles represent nodes where cell trajectory branches into several different outcomes. Numbers inside each node are for reference purposes only, and do not hold other meaning.

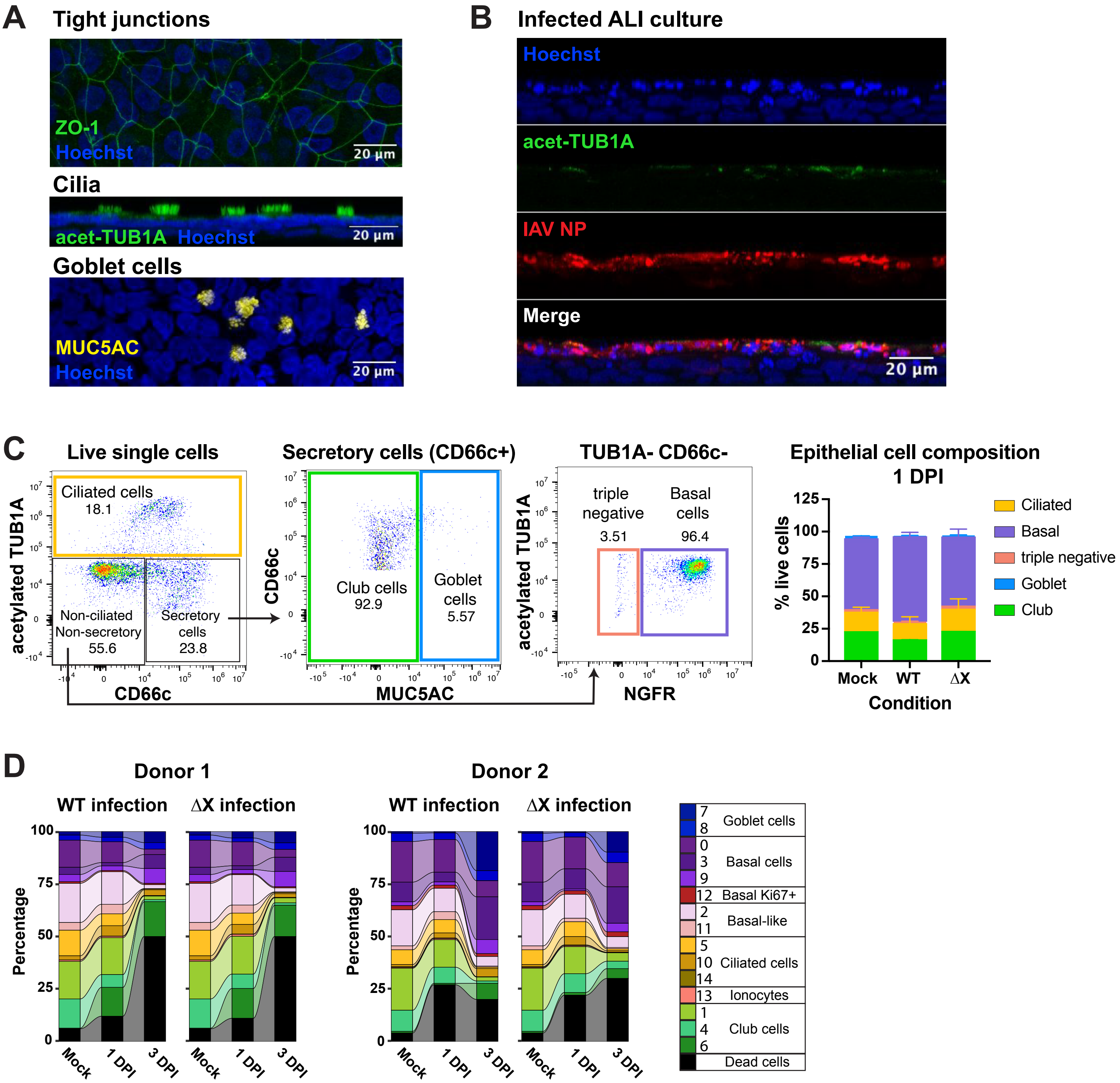
A
Tight junctions
ZO-1
Hoechst
20 µm
Cilia
acet-TUB1A Hoechst
20 µm
Goblet cells
MUC5AC
Hoechst
20 µm
B
Infected ALI culture
Hoechst
acet-TUB1A
IAV NP
Merge
20 µm
C
Live single cells
Ciliated cells
18.1
Non-ciliated Non-secretory
55.6
Secretory cells
23.8
acetylated TUB1A
CD66c
Secretory cells (CD66c+)
Club cells
92.9
Goblet cells
5.57
CD66c
MUC5AC
TUB1A- CD66c-
triple negative
3.51
Basal cells
96.4
acetylated TUB1A
NGFR
Epithelial cell composition
1 DPI
% live cells
Ciliated
Basal
triple negative
Goblet
Club
Mock
WT
ΔX
Condition
D
Donor 1
Donor 2
WT infection
ΔX infection
Percentage
Mock
1 DPI
3 DPI
7
8
Goblet cells
0
3
9
Basal cells
12 Basal Ki67+
2
11
Basal-like
5
10
14
Ciliated cells
13 Ionocytes
1
4
6
Club cells
Dead cells

**Figure S2. Influenza A viral infection alters cell type composition in the airway epithelium in a PA-X-independent manner.** ALI cultures were infected and samples collected as described in Figure 1D. scRNAseq data from Perth H3N2-infected ALI cultures were analyzed to identify groups of cells with similar expression patterns and to assign cell type (n = 2, Donors 1 and 2). A) Human bronchial epithelial cells were cultured at ALI for 3-4 weeks and stained for tight junctions (ZO-1, top), cilia (acetylated TUB1A, middle, sagittal section), and the mucus component and goblet cell marker mucin 5AC (MUC5AC, bottom) and nuclear stain (Hoescht, all) to confirm proper differentiation of the cells and formation of the pseudostratified epithelium. B) Immunofluorescence image of infected ALI culture, with staining of influenza A virus nucleoprotein (NP), cilia (acetylated TUB1A), and nuclear stain (Hoechst) at 2 days post infection. C) Cells were collected for flow cytometric validation of cell types at 1 day post infection (DPI). (Left) Gating strategy for epithelial cell types in ALI cultures after gating for live, single cells. We termed cells negative for ciliated, secretory, and basal markers "triple-negative" cells. They likely represent the cells in cluster 13 (Figure 1E). (Right) Stacked bar plot of average epithelial cell type composition from two different donors (Donors 4, 5). D) Alluvial plot showing the change in proportions of cells per cluster and cell type throughout infection with Perth H3N2 WT and Δ X. The two donors are plotted separately. The percentage of dead cells was calculated based on cell viability measurements using trypan blue carried out prior to dead cell depletion for scRNAseq library preparation.

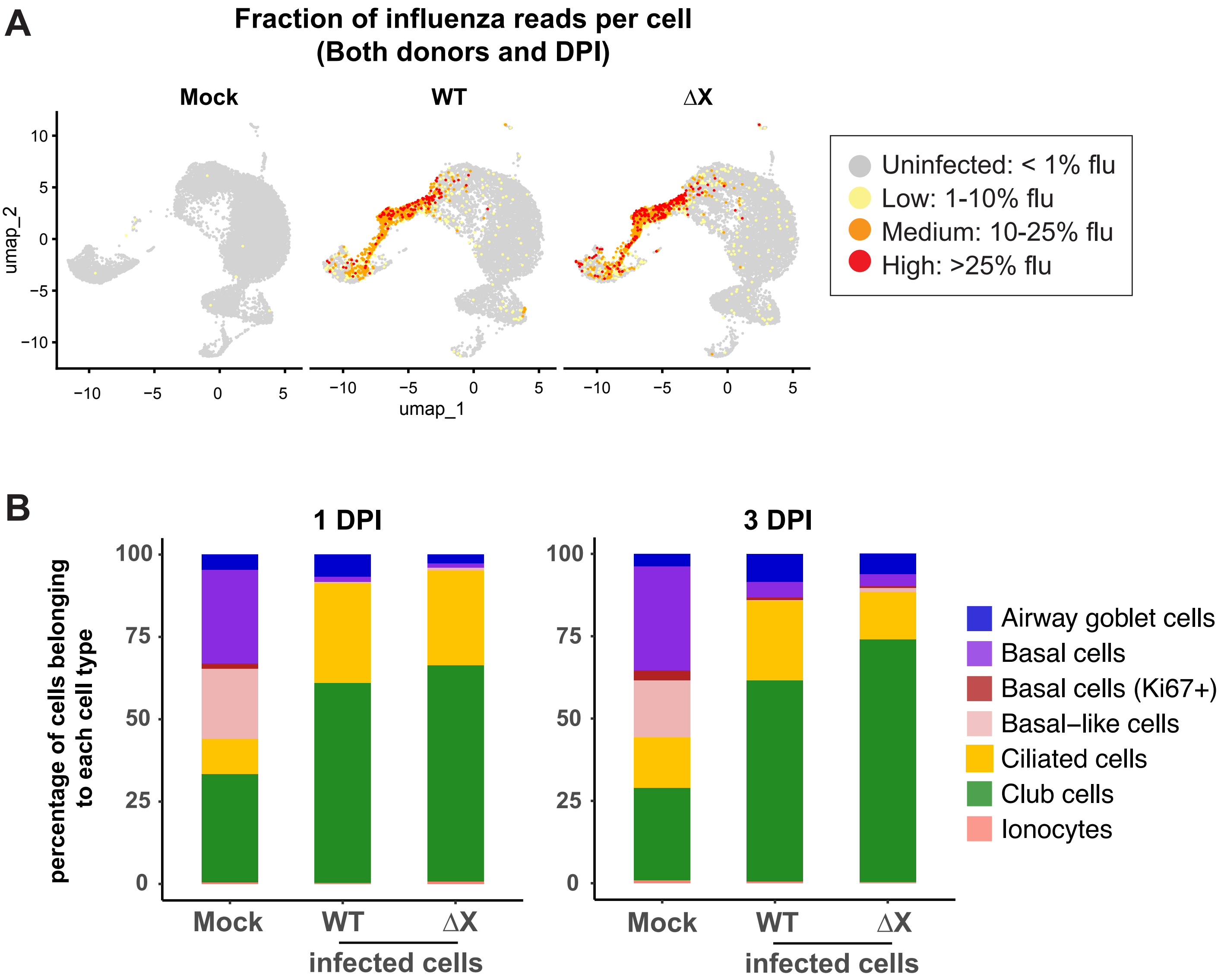


**Figure S3. PA-X does not alter infection parameters at the single cell level.** ALI cultures were infected and samples collected as described in Figure 1D. The scRNAseq data were analyzed to identify infected cells (n = 2, Donors 1 and 2). A) Integrated UMAP of mock, Perth H3N2 WT and Perth H3N2 ΔX infected ALI cultures colored by the percent of reads that mapped to influenza A viral genes in each cell. B) Bar graph showing the percentage of infected cells that belong to each epithelial cell type vs. the composition of mock-infected cultures. Cells were deemed infected if at least 1% reads mapped to the influenza transcriptome. The two donors were aggregated for the analysis in all panels. The two timepoints (1 and 3 DPI) were aggregated for panel A.

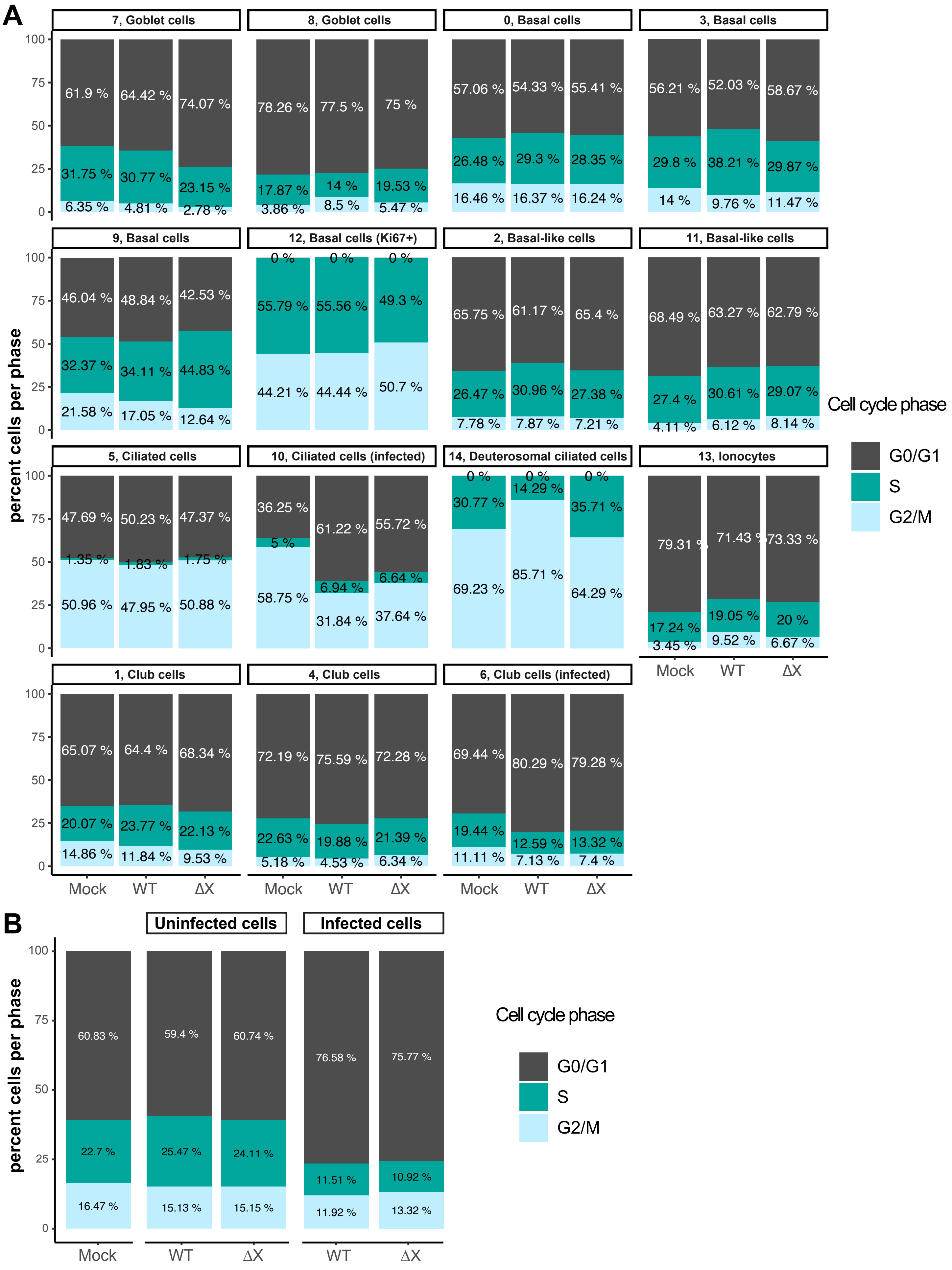
A
7, Goblet cells
61.9 % 64.42 % 74.07 %
31.75 % 30.77 % 23.15 %
6.35 % 4.81 % 2.78 %
8, Goblet cells
78.26 % 77.5 % 75 %
17.87 % 14 % 19.53 %
3.86 % 8.5 % 5.47 %
0, Basal cells
57.06 % 54.33 % 55.41 %
26.48 % 29.3 % 28.35 %
16.46 % 16.37 % 16.24 %
3, Basal cells
56.21 % 52.03 % 58.67 %
29.8 % 38.21 % 29.87 %
14 % 9.76 % 11.47 %
9, Basal cells
46.04 % 48.84 % 42.53 %
32.37 % 34.11 % 44.83 %
21.58 % 17.05 % 12.64 %
12, Basal cells (Ki67+)
0 % 0 % 0 %
55.79 % 55.56 % 49.3 %
44.21 % 44.44 % 50.7 %
2, Basal-like cells
65.75 % 61.17 % 65.4 %
26.47 % 30.96 % 27.38 %
7.78 % 7.87 % 7.21 %
11, Basal-like cells
68.49 % 63.27 % 62.79 %
27.4 % 30.61 % 29.07 %
4.11 % 6.12 % 8.14 %
5, Ciliated cells
47.69 % 50.23 % 47.37 %
1.35 % 1.83 % 1.75 %
50.96 % 47.95 % 50.88 %
10, Ciliated cells (infected)
36.25 % 61.22 % 55.72 %
5 % 6.94 % 6.64 %
58.75 % 31.84 % 37.64 %
14, Deuterosomal ciliated cells
0 % 0 % 0 %
30.77 % 14.29 % 35.71 %
69.23 % 85.71 % 64.29 %
13, Ionocytes
79.31 % 71.43 % 73.33 %
17.24 % 19.05 % 20 %
3.45 % 9.52 % 6.67 %
1, Club cells
65.07 % 64.4 % 68.34 %
20.07 % 23.77 % 22.13 %
14.86 % 11.84 % 9.53 %
4, Club cells
72.19 % 75.59 % 72.28 %
22.63 % 19.88 % 21.39 %
5.18 % 4.53 % 6.34 %
6, Club cells (infected)
69.44 % 80.29 % 79.28 %
19.44 % 12.59 % 13.32 %
11.11 % 7.13 % 7.4 %
percent cells per phase
0 25 50 75 100
Mock WT ΔX
Cell cycle phase
G0/G1
S
G2/M
B
Uninfected cells
Infected cells
60.83 % 59.4 % 60.74 % 76.58 % 75.77 %
22.7 % 25.47 % 24.11 % 11.51 % 10.92 %
16.47 % 15.13 % 15.15 % 11.92 % 13.32 %
percent cells per phase
0 25 50 75 100
Mock WT ΔX WT ΔX
Cell cycle phase
G0/G1
S
G2/M

**Figure S4. Influenza A virus infected cells arrest in G0/G1, while bystander cells do not.** Analysis of scRNAseq data from ALI cultures infected as shown in Figure 1D (n = 2, Donors 1 and 2). A-B) Bar graphs showing analysis of cell cycle phase using Seurat at 1 day post infection (DPI), shown according to A) cluster and assigned cell type or B) infection status. Infected cells = influenza A virus genes ≥1% of total reads; uninfected cells = influenza A virus genes < 1% of total reads.

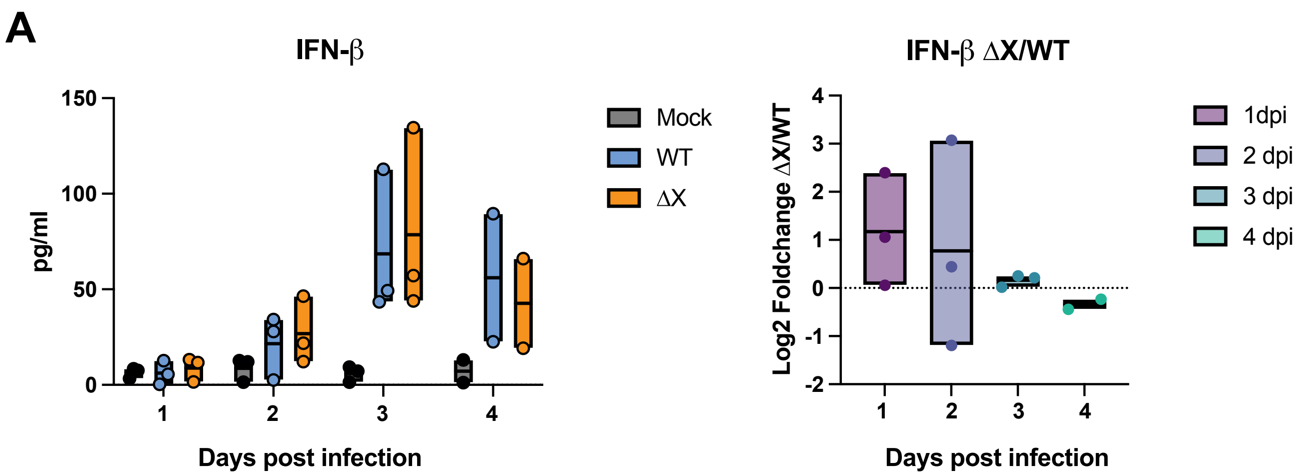


**Figure S5. PA-X does not alter IFN-β secretion during infection.** ALI cultures were infected with Perth H3N2 WT or ΔX or mock infected at MOI 0.1. Basal media was collected every day after infection and IFN-β protein levels were tested using ELISA. A) IFN-β protein concentration in basal media. C) Log2 fold-change of IFN- secreted by ΔX-infected ALIs vs. WT-infected (n = 3 experiments in donor 1).

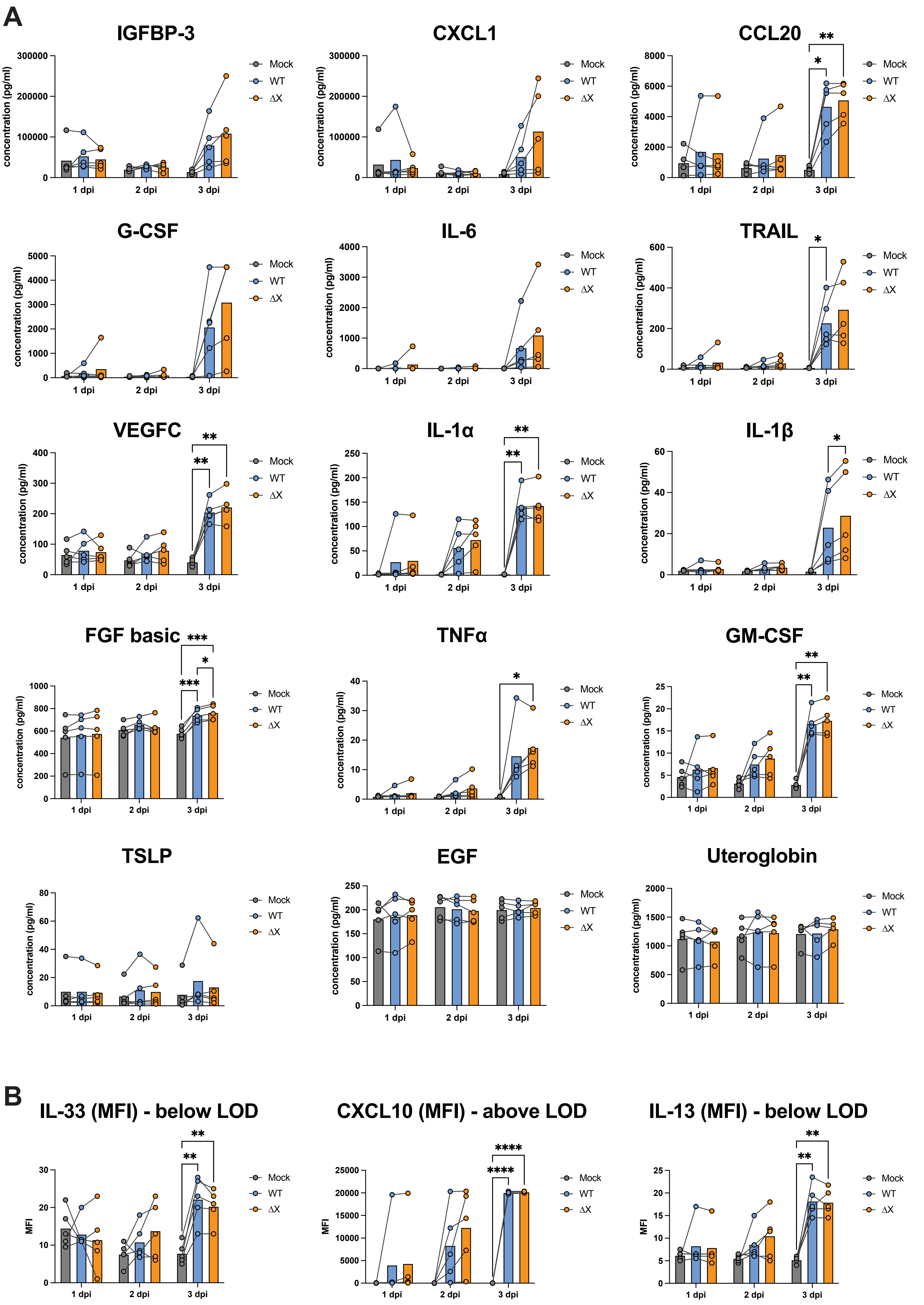
A
IGFBP-3
CXCL1
CCL20
G-CSF
IL-6
TRAIL
VEGFC
IL-1α
IL-1β
FGF basic
TNFα
GM-CSF
TSLP
EGF
Uteroglobin
concentration (pg/ml)
1 dpi
2 dpi
3 dpi
Mock
WT
ΔX
B
IL-33 (MFI) - below LOD
CXCL10 (MFI) - above LOD
IL-13 (MFI) - below LOD
MFI

**Figure S6. PA-X limits secretion of pro-inflammatory and lung injury-associated cytokines**
A-B) Luminex data using basal conditioned media from ALI cultures infected as shown in Figure 1D (n = 5, 5 separate donors, Donors 1-5). B) Bar graphs showing concentrations of cytokines throughout infection time course. Comparisons between conditions are shown (analyzed using a two-way ANOVA, followed by Šídák's multiple comparisons test. ns = $p > 0.05$, *= $p \leq 0.05$, **= $p \leq 0.01$, ***= $p \leq 0.001$.) B) For targets that were outside the limit of detection (LOD) of the standard curve, median fluorescence intensity is plotted in bar graphs. Comparisons between conditions are shown (analyzed using a two-way ANOVA. ns = $p > 0.05$, *= $p \leq 0.05$, **= $p \leq 0.01$, ***= $p \leq 0.001$.)

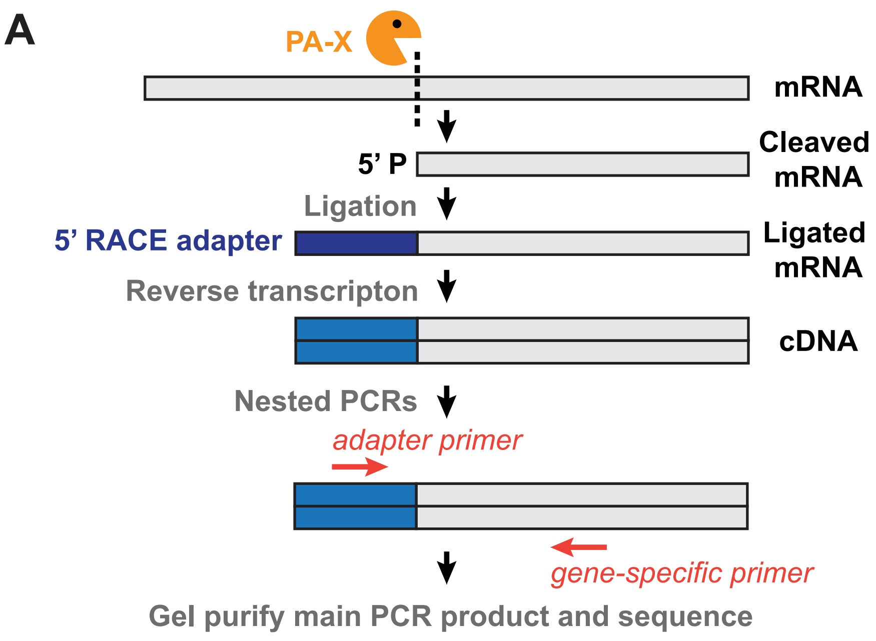


**B**

| Gene name | # predicted cut sites |
|---|---|
| *CALR* | 4 |
| *CANX* | 5 |
| *HSP90AA1* | 4 |
| *HSP90AB1* | 7 |
| *PSME1* | 1 |
| *PSME2* | 1 |

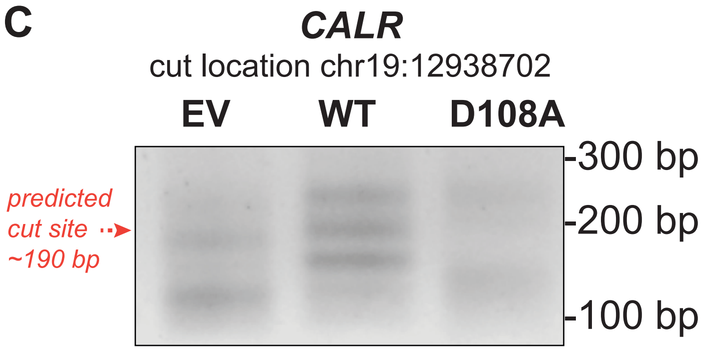


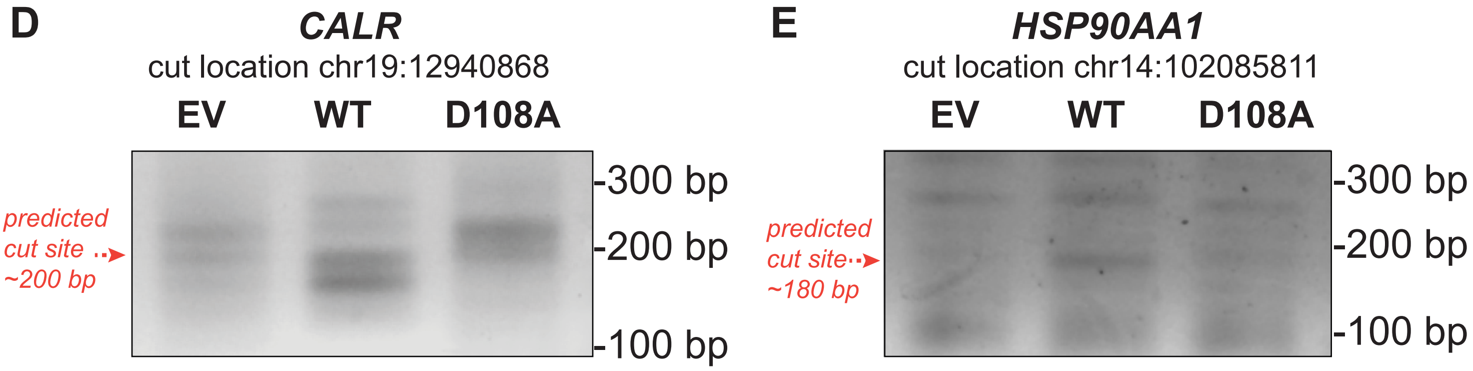


**Figure S7. PA-X directly cleaves antigen processing and presentation genes.** A) Schematic of the 5’ RACE workflow. 5’ P, 5’ phosphate. B) Table of cut sites detected by 5’ RACE-seq analysis in Gaucherand et al. 2023 in the significantly downregulated genes listed in Figure 4B. Arrows indicated the sites that were selected for experimental validation. C-E) XRN1 inducible knockdown HEK 293T cells were transfected with empty vector, WT Perth H3N2 PA-X, or catalytically inactive (D108A) Perth H3N2 PA-X for 24 hours before RNA extraction. 5’ RACE was then performed using primers downstream of cut sites identified in the 5’ RACE-seq in (C-D) CALR and (E) HSP90AA1. Arrows indicate the position of PCR fragments that correspond to the expected cut sites. The DNA bands were purified and sequenced to confirm their identities. Additional bands in control samples are derived from RNA fragments present in the cell but unrelated to PA-X activity.

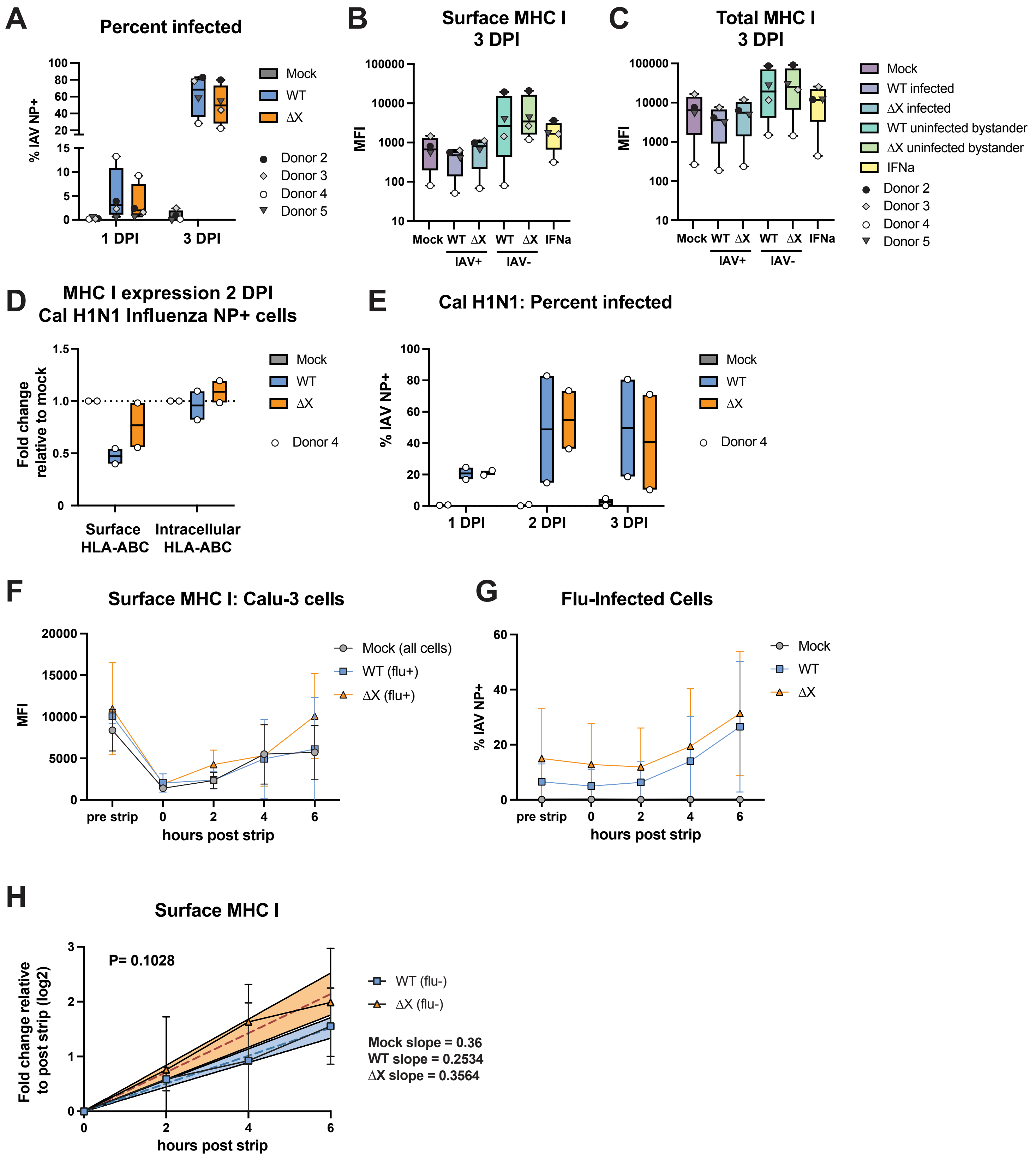
A
Percent infected
% IAV NP+
Mock
WT
ΔX
Donor 2
Donor 3
Donor 4
Donor 5
1 DPI
3 DPI
B
Surface MHC I
3 DPI
MFI
Mock WT ΔX WT ΔX IFNa
IAV+
IAV-
C
Total MHC I
3 DPI
MFI
Mock
WT infected
ΔX infected
WT uninfected bystander
ΔX uninfected bystander
IFNa
D
MHC I expression 2 DPI
Cal H1N1 Influenza NP+ cells
Fold change relative to mock
Surface HLA-ABC
Intracellular HLA-ABC
E
Cal H1N1: Percent infected
% IAV NP+
1 DPI
2 DPI
3 DPI
F
Surface MHC I: Calu-3 cells
MFI
Mock (all cells)
WT (flu+)
ΔX (flu+)
pre strip
hours post strip
G
Flu-Infected Cells
% IAV NP+
H
Surface MHC I
P= 0.1028
Fold change relative to post strip (log2)
WT (flu-)
ΔX (flu-)
Mock slope = 0.36
WT slope = 0.2534
ΔX slope = 0.3564
hours post strip

**Figure S8. MHC I levels during infection.** A-C) ALI cultures were infected with Perth H3N2 WT or ΔX virus, or mock-infected at MOI 0.1. Fixed cells were stained with an anti-HLA-ABC antibody prior to permeabilization to detect the surface exposed portion of MHC-I. The cells were then permeabilized and stained for internal HLA-ABC, using the same antibody coupled with a different fluorophore. Cells were also stained for influenza A virus nucleoprotein (NP). IAV+ = influenza A virus NP-positive cells. A) Infected cells NP+ cells at 1 and 3 days post infection (DPI) as a percent of total live cells. B) Median fluorescence intensity of surface HLA-ABC. C) Median fluorescence intensity of intracellular HLA-ABC. D-E) ALI cultures were infected with Cal H1N1 WT or ΔX virus, or mock-infected at MOI 0.1. Cells were stained as described for A-C. D) HLA-ABC median fluorescence intensity at 2 DPI on NP+ infected cells from flow cytometry analysis plotted as fold-change relative to mock-infected ALI cultures. E) Infected cells NP+ cells at 1-3 DPI as a percent of total live cells. F-H) Calu-3 cells were mock-infected or infected with Perth H3N2 WT or ΔX at an MOI of 0.1 for 18 hours. Surface MHC I was stripped using a citric acid buffer, then cells were collected at multiple timepoints over the course of 6 hours. F) Median fluorescence intensity of MHC I on the surface of Calu-3 cells during this experiment. G) Percent of influenza NP+ cells. H) Log2 fold change of surface MHC I median fluorescence intensity over a 6-hour time course in influenza A virus NP- bystander cells in WT and ΔX-infected cultures. Linear regression was performed for each condition, and the p value was calculated to compare Perth H3N2 WT and ΔX NP- bystander cells. Dashed lines represent the line of best fit, and the shaded regions indicate the 95% confidence interval (see Figure 4F for equivalent plot for mock-infected cells).
